# Snorkel-tag Based Affinity Chromatography for Recombinant Extracellular Vesicle Purification

**DOI:** 10.1101/2024.04.12.589209

**Authors:** Madhusudhan Reddy Bobbili, André Görgen, Yan Yan, Stefan Vogt, Dhanu Gupta, Giulia Corso, Samir Barbaria, Carolina Patrioli, Sylvia Weilner, Marianne Pultar, Jaroslaw Jacak, Matthias Hackl, Markus Schosserer, Regina Grillari, Jorgen Kjems, Samir EL Andaloussi, Johannes Grillari

## Abstract

Extracellular vesicles (EVs) are lipid nanoparticles and play an important role in cell-cell communications, making them potential therapeutic agents and allowing to engineer for targeted drug delivery. The expanding applications of EVs in next generation medicine are still limited by existing tools for scaling standardized EV production, single EV tracing and analytics, and thus provide only a snapshot of tissue-specific EV cargo information. Here, we present CD81, an EV surface marker protein, genetically fused to series of tags with additional transmembrane domain to be displayed on the EV surface, which we term Snorkel-tag. This system enables to affinity purify EVs from complex matrices in a non-destructive form. In future applications, this strategy will allow generating transgenic animals to enable tracing and analyzing EVs, and their cargo in physiological and pathophysiological set-ups, and facilitate the development of EV based diagnostic tools in murine models which can be translated to humans.

## Introduction

Extracellular vesicles (EVs <200 nm) are lipid bilayer nanoparticles secreted by all cells from prokaryotes to eukaryotes. EVs play a key role in intercellular communication, transferring bioactive molecules to recipient cells or tissues thus modulating a plethora of functions. EVs are heterogenous in nature with sizes ranging from 30 – 1000 nm and are broadly classified into two classes, exosomes that are < 150 nm in diameter formed within multivesicular bodies (MVBs) and released upon fusion to plasma membrane, and the ectosomes >150 nm released by plasma membrane budding^1,2^. Exosomes and ectosomes can be identified by the presence of multiple cellular components such as tumor susceptibility gene 101 protein (TSG101), Alix, syndecan-syntenin complexes and tetraspanin family (CD63, CD9, and CD81), involved in their biogenesis^3–5^. During the last two decades, studies on EVs have shown that tetraspanins CD9, CD63 and CD81 are highly enriched broadly on the surface of all classes of EVs^6–8^. Recent studies report, EVs enriched for CD63 correspond to endosome-derived exosomes, while plasma membrane-derived ectosomes mostly bear CD9^9,10^. CD81 appears to be highly enriched in both, exosomes and to some extent in ectosomes^10^.

There is accumulating evidence showing that mesenchymal stem cell (MSC)-derived EVs have powerful therapeutic effects when derived of parent MSCs with low immunogenicity^11^. In addition, their ability to cross blood-tissue barriers, offers unique platform to engineer EVs for targeted drug-delivery^12,13^. Due to their diverse sources and respective changes in their cargo content including nucleic acids, lipids, glycans, proteins, and metabolites in various physiological and pathophysiological conditions, EVs offer perspectives for multi-analyte biomarker measurements. However, our understanding of EV biogenesis, cargo loading, uptake and regulatory mechanisms involved come mostly from in vitro studies. The lack of understanding EV biogenesis, composition and function in vivo remain a critical challenge to exploit EVs to their full potential for development of EV-based biomarker and therapeutics^14^. In all these applications, tools to unambiguously affinity-purify EVs in a non-destructive manner from cell culture supernatants or from biological samples including tissues or biofluids such as serum, plasma, CSF or urine are still lacking, despite a variety of enrichment strategies available. Therefore, here we report on a versatile and broadly applicable pre-clinical tool for generating EVs carrying a CD81 fusion protein that displays a series of tags for labeling and non-destructive cleavage from affinity columns on the EV surface.

## Results

### Design and validation of Snorkel-tag

To design a versatile tag for broad applications of EVs, we first selected an EV marker protein that should be ubiquitous to all EVs. For this reason, we chose the tetraspanin family members CD63 and CD81 as highly enriched in all classes of EVs^10^. Second, the tag should be displayed on the EV surface and be versatile, in order to enable affinity purification and tracking of EVs. As tetraspanins are multipass membrane proteins with N- and C-termini located intravesicularly, the tags must be either inserted into one of the extracellular loops of tetraspanins which can produce unintended biochemical changes leading to protein misfolding, instability, altered post-translational modifications, and aberrant enrichment in EV membranes leading to functional changes^15–17^ or omitting/adding transmembrane spanning regions to have one of the termini displayed on the surface can result in misfolding or wrong subcellular localization^18,19^. These considerations led us to generate a series of genetic constructs with CD63 and CD81 fusion proteins either lacking one transmembrane domain or by adding a fifth transmembrane domain derived from mouse beta-type platelet-derived growth factor PDGFRB^20^ (Uniprot accession <u>P05622</u>) to display the respective N- or C-terminus to the outside of the EVs (Fig S1).

From these constructs, we selected the correctly membrane localizing fusion of CD81 (Fig S2) to a fifth transmembrane domain at its C-terminus with an additional extravesicular domain consisting of an HA-tag, a PreScission protease cleavage site for non-destructive elution from affinity columns, a CLIP-tag and a FLAG-tag for further staining with anti-FLAG antibodies or covalently label with CLIP substrates. Additionally, flexible linkers [(G4S)3] were introduced on either side of the PreScission protease to avoid steric hindrance of antibody binding and protease cleavage during affinity purification. This series of tags together were termed Snorkel-tag^15^ (scheme in Supplementary note A, Fig 1A). In this study, as a proof-of-principle, we preferred CD81 over CD63 due to its high enrichment in small EVs, while obviously any other tetraspanin molecule of EVs might be amenable to fusion with the Snorkel-tag.

**Figure 1:**
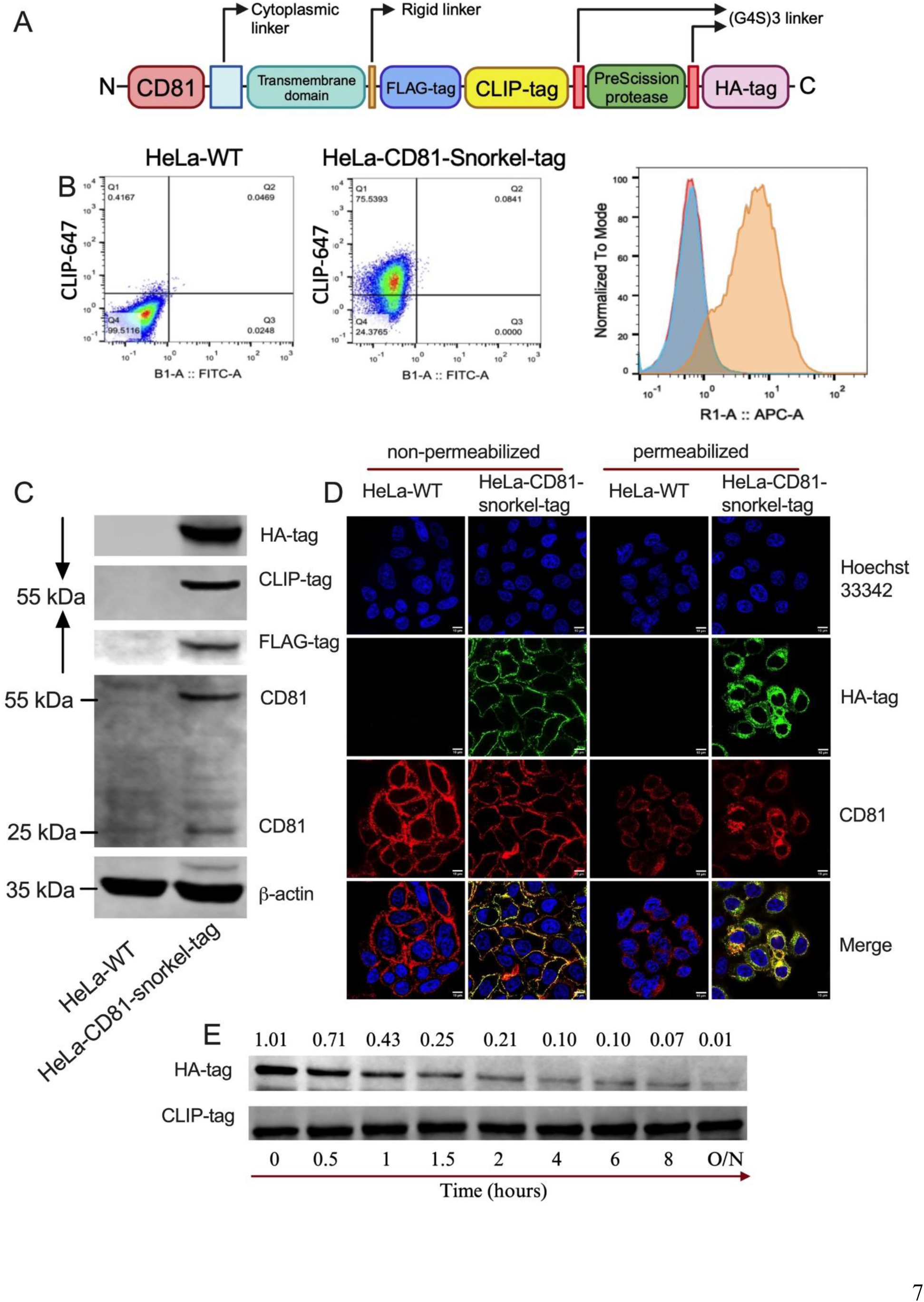
Expression of CD81-Snorkel-tag in HeLa cells results in accessibility of all parts of the Snorkel-tag. **A,** Schematic illustration of snorkel-tag fused to C-termini of CD81. **B,** Flow cytometric analysis of HeLa-WT and HeLa-CD81 snorkel-tag stable cells stained with CLIP membrane impermeable substrate CLIP-647. **C,** Western blot of HA-tag, FLAG-tag, CLIP-tag, CD81 and beta-actin for WT and CD81-snorkel-tag cell lines. Beta-actin was used as loading control. **D,** Fluorescent images of fixed WT and CD81-snorkel-tag cells. Right two panels: Immunofluorescence (IF) for CD81 and HA-tag in non-permeabilized HeLa cells, Left two panels: IF for CD81 and HA-tag in permeabilized HeLa cells. Counterstaining with Hoechst 33342 for DNA. **E,** Western Blot of HeLa cells expressing CD81-snorkel-tag treated with PreScission protease for indicated time periods mentioned above. The blots were probed with antibodies against HA-tag and CLIP-tag. The mean intensity of the bands was quantified and the ratio of the HA-tag vs. CLIP-tag is shown.

To produce recombinant Snorkel-tagged EVs, we stably expressed the CD81-snorkel-tag construct in HeLa and in telomerized mesenchymal stroma cells (WJ-MSC/TERT273) by the pLVX lentiviral transduction system. Stable cells were characterized for detectability of all snorkel-tag components by flow cytometry for CLIP-tag using membrane impermeable fluorescent CLIP substrate (Fig 1B) and immunoblotting for HA-tag, FLAG-tag and CLIP-tag (Fig 1C). Confocal microscopy confirmed localization of CD81-snorkel-tag to the cell membrane on non-permeabilized as well as intracellularly in permeabilized cells (Fig 1D). Cleavability of the PreScission protease cleavage site was assessed and optimized by enzymatic digestion of whole cell lysates (Fig 1E). Taken together, these experiments confirm the expression of Snorkel-tag on the plasma membrane and its functionality and versatility for affinity purification and labelling.

### Generation and characterization of EVs for Snorkel-tag

To achieve the ultimate goal of non-destructive affinity purification of EVs, we enriched EVs from HeLa-CD81-snorkel-tag and HeLa-WT cells as controls from conditioned media by tangential flow filtration (TFF). EV preparations were characterized for their concentrations and diameter by nanoparticle tracking analysis (NTA) (Fig 2A, B). TFF isolated EV preparations were enriched for Snorkel-tag components (HA-tag, CLIP-tag and FLAG-tag), classical EV-specific markers (Alix, Syntenin-1) and depleted from other intracellular proteins such as Calnexin, an endoplasmic reticulum specific protein (Fig 2C). Transmission electron microscopy (TEM) validated size and morphology of EVs in the expected range, as well as presence of CD81 and the HA-tag by immunogold labelling (Fig 2D). We next evaluated if the EV protein surface signature would change due to CD81 snorkel-tag overexpression using multiplex bead-based flow cytometry (MBFCM)^21^. EVs from both WT and CD81 Snorkel-tag overexpressing cells showed robust expression of abundant surface markers like tetraspanins, CD44, MCSP-1 and CD146 with pan tetraspanin detection. As negative control, we used anti-HA antibody and Dylight-649 secondary antibody for detection, and did not detect any background for WT EVs as expected, while we observed clear signals on CD81 snorkel-tag enriched EVs for tetraspanins, CD44, MCSP-1 and CD146. This indicates that overexpression of CD81 Snorkel-tag did not alter the EV protein surface signature (Fig 2E). Together, these results provide evidence that the snorkel-tag fused to the C-terminus of CD81 is well displayed on the surface of the EV membranes.

**Figure 2:**
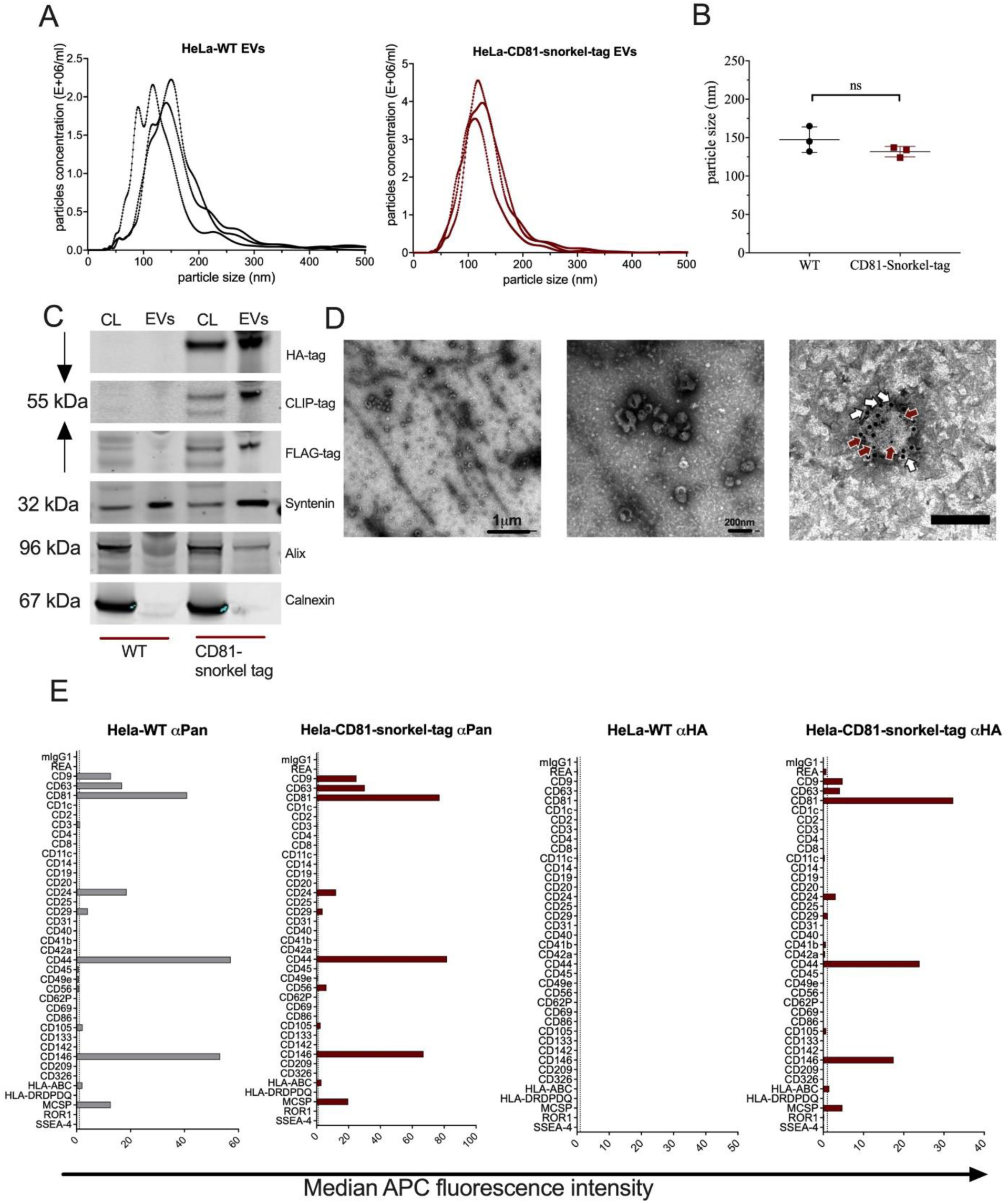
Snorkel-tagged EVs isolated by tangential flow filtration (TFF) show similar characteristics as wild type EVs. **A** and **B,** NTA analysis of TFF isolated EVs from HeLa-WT and HeLa-CD81-snorkel-tag for particle concentration and size. **C,** Western blot of HeLa-WT and HeLa-CD81-snorkel-tag cell lysates and EV lysates for snorkel-tag epitopes and EV specific markers. **D,** Transmission electron microscopy (TEM) images for HeLa-WT EVs and HeLa-CD81-snorkel-tag carrying EVs. Overview image (Left, scale bar 1 µm) and close-up (right, scale bar 200 nm). White arrows label CD81 (anti-mouse antibody tagged to10 nm gold particle) and red arrows label Snorkel-tag (anti-rabbit antibody tagged to 4 nm gold particle) (scale bar: 100 nm). **E,** Multiplex bead-based flow cytometry assay for detection of EV surface protein signature by pan tetraspanin detection antibodies and anti-HA antibody.

### Snorkel-tag based Extracellular Vesicle Affinity Chromatography (STEVAC)

To evaluate the purification of our recombinant EVs, we enriched them from HeLa-CD81-Snorkel-tag and, as control, from HeLa-WT conditioned media by ultrafiltration using 100 MWCO Amicon filters and quantified concentration and size by NTA (Fig S3A, B). Next, we performed StEVAC on the enriched EVs. In brief, EVs were incubated with anti-HA magnetic beads overnight at 4° C. Post incubation, beads were washed to remove uncaptured EVs and non-specifically bound components. The captured EVs on the beads were then incubated with PreScission protease overnight at 4° C for on-column cleavage (Fig 3A, Supplementary notes B) and non-destructive elution of EVs, as opposed to using EV destroying high salt or low pH buffers necessary to disrupt antigen-antibody binding as usually used for elution in affinity chromatography.

**Figure 3:**
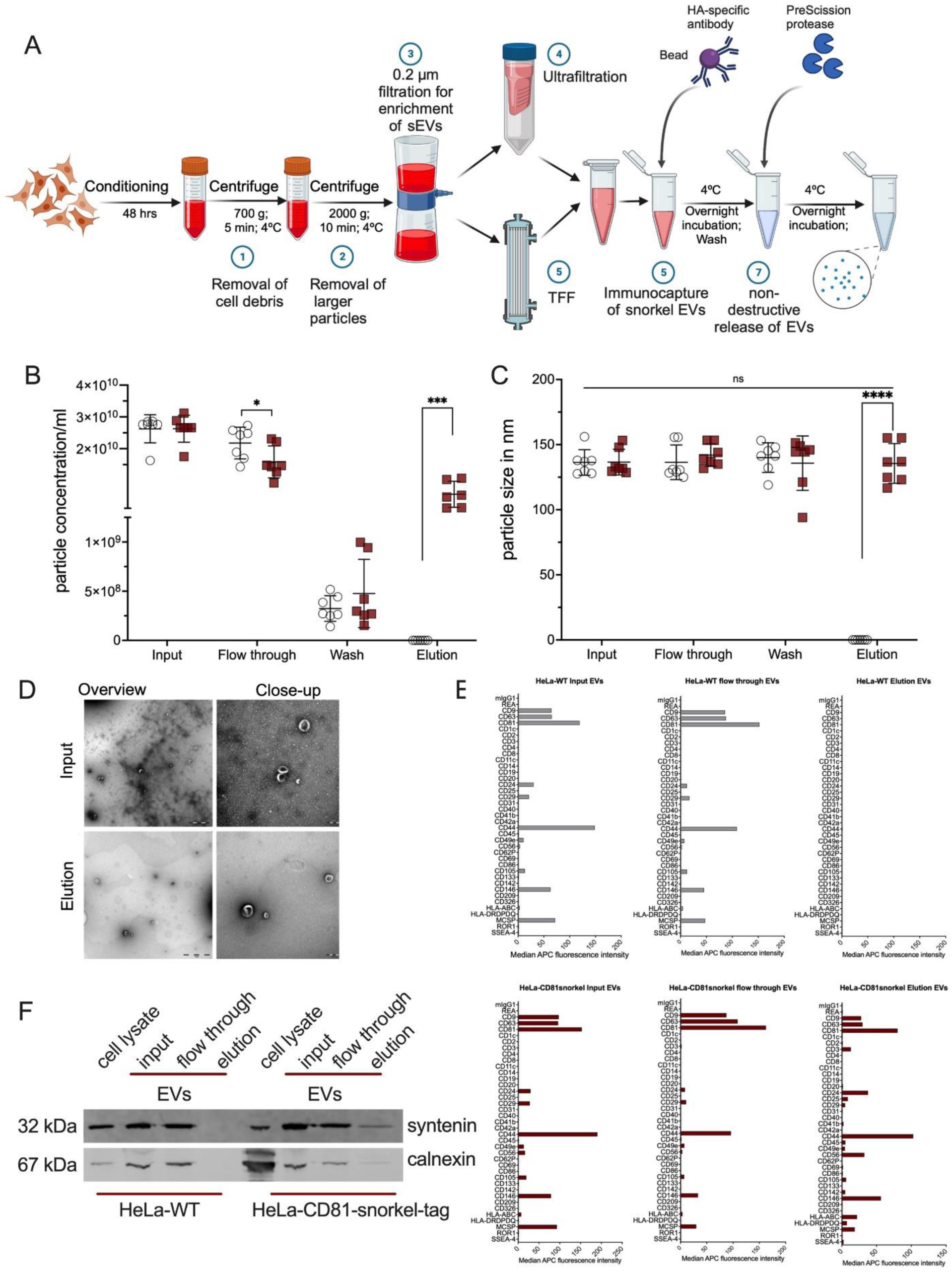
Snorkel-tag based Extracellular Vesicle Affinity Purification. **A,** Schematic illustration of experimental setup for StEVAC method. **B,** Nanoparticle tracking analysis (NTA) counts and **C**, particle diameter of purified EVs eluted after incubation with PreScission protease overnight and a following wash step (n=7). **D,** Transmission electron microscopic examination for size and morphology of the input and eluted EVs (left panel with overview image, scale bar 1µm; right panel with close-up image, scale bar 200nm). **E,** Multiplex bead-based flow cytometry assay to evaluate the StEVAC method. Assay results for HeLa-WT (top) and HeLa-CD81-Snorkel-tag (bottom); input, flow through (unbound EVs) and elution. (n=3). **F,** Western blots for the EV-associated protein syntenin and non-EV marker calnexin (ER specific) in the elutes. 1-way ANOVA was applied on raw values; nsP > 0.05, **P < 0.01, ***P < 0.001.

StEVAC was performed in 3-7 independent replicates using ∼2.5 x 10^10^ particles/mL as input and wild type EVs as control. EVs in eluates were quantified by NTA showing a recovery of around 30% of input^8,9^ (Fig 3B). No differences in size profiles of unbound or eluted particles were observed (Fig 3C). Next, we performed a detailed characterization of StEVAC purified Snorkel-tag EVs. EVs show the expected size in TEM, where we also observed a strong reduction of background signals (Fig 3D). MBFCM on flow-throughs and elutes showed the same surface protein pattern as for input EVs, while WT elutes served as negative controls (Fig 3E), indicating the specificity of the method. Immunoblotting of the various fractions of StEVAC show de-richment of calnexin as a marker for cytoplasmic contaminations (Fig 3F).

### Confirming specificity of StEVAC

We next confirmed the specificity of StEVAC by pre-blocking anti-HA magnetic beads with an excess of HA peptide before incubation with CD81 Snorkel-tag EVs. NTA results demonstrate that pre-blocking HA matrix with HA peptide indeed resulted in no recovery in the elutes (Fig 4A).

**Figure 4:**
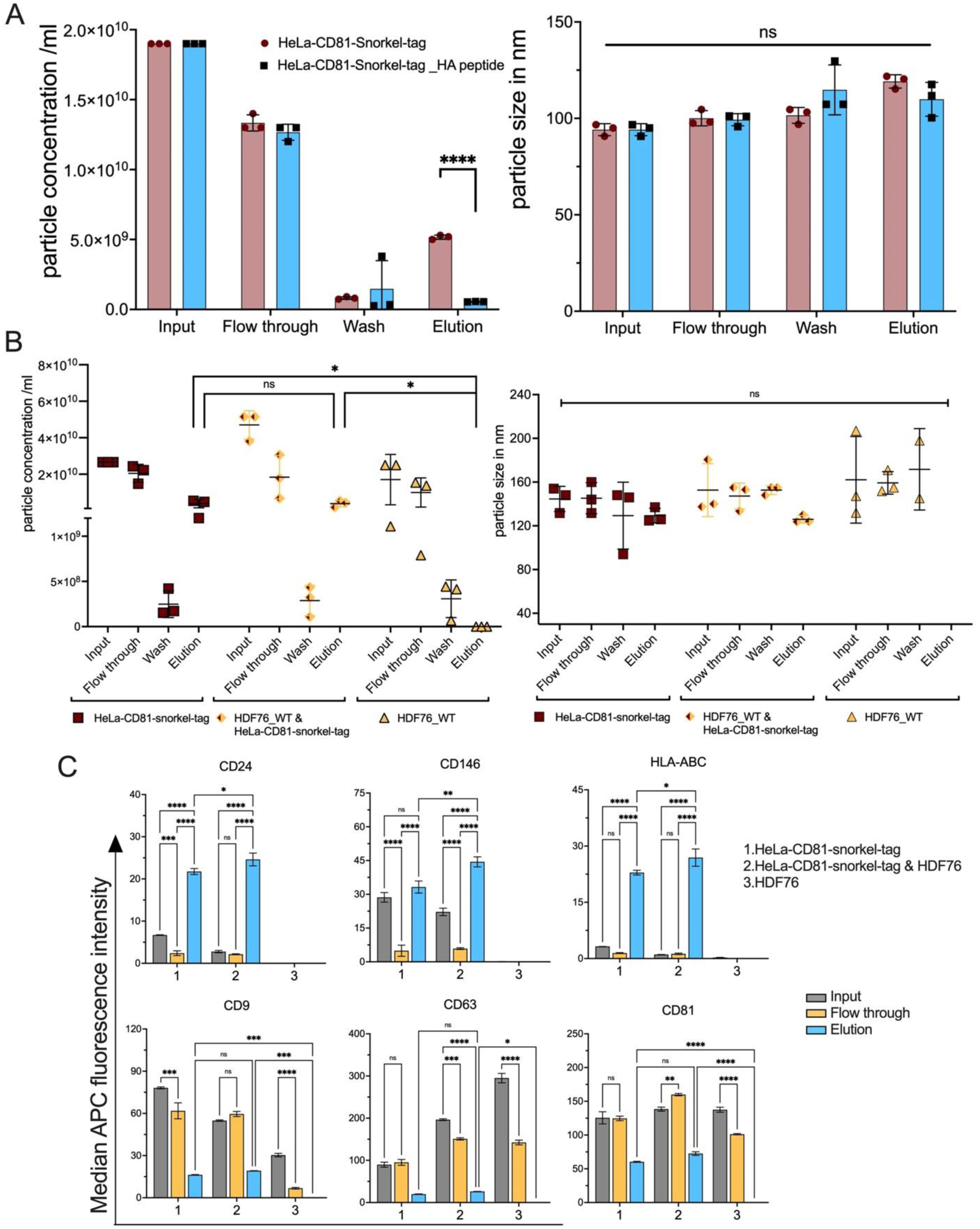
StEVAC specifically isolates snorkel-tagged EVs out of complex mixtures. **A,** Pre-binding of anti-HA matrix with HA peptide shows no binding of CD81-Snorkel-tag EVs henceforth no EVs in elution. Nanoparticle analysis (NTA) shows significant differences in elutions between anti-HA matrix pre-bound with HA peptide and free paratope (n=3). **B,** Nanoparticle tracking analysis (NTA) counts of purified EVs from HeLa-CD81-Snorkel-tag, HDF76 and mix of both, eluted after incubation with PreScission protease treatment overnight and a following wash step (n=3). No significant differences were observed in size distribution between flowthrough, washes and elution compared to input between CD81-Snorkel-tag EVs alone or mixed HDF76 EVs. **C,** Multiplex bead-based assay results for input, flowthrough and elution of EVs from HeLa-CD81-Snorkel-tag, HDF76 and HeLa-CD81-Snorkel-tag mixed with HDF76. CD24, CD146 & HLA-ABC are enriched only in the elutions of HeLa-CD81-Snorkel-tag EVs (D; top row), additionally, tetraspanins CD9, CD63 & CD81 are enriched in all the elutions except in HDF76. Particle size quantification revealed no significant size difference during the isolation process (n=3). One-way ANOVA multiple comparison test was used for analysis of (A); 2-way ANOVA multiple comparison test was used for analysis of (B) and (C). Data are shown as mean+_SD, *P < 0.05, **P < 0.01, ***P < 0.001.

To test, if Snorkel-tag EVs can be purified from EV mixtures, we combined HeLa-CD81-Snorkel-tag EVs with primary human dermal fibroblast (HDF76) untagged EVs and performed StEVAC. We observed concentration of particles in the elute from mixed EV inputs were similar to the CD81 Snorkel-tag EVs elutes (Fig 4B). Furthermore, we quantified diameter and EV protein signatures by multiplex bead-based flow cytometry (MBFCM) before and after StEVAC. None of the 37 surface proteins included in the MBFCM panel were specific to fibroblast EVs, therefore, we had to rely on quantitating EV numbers and surface marker signal intensities of HeLa specific proteins CD24, CD146 and HLA-ABC to estimate enrichment (Fig 4C, Fig S4A-C).

### StEVAC method enables purification of Snorkel-tag EVs from complex matrices

To showcase the potential of StEVAC, we then asked whether Snorkel-tag EVs can be purified from more complex matrices such as human platelet derived EVs and plasma for removal of contaminating EVs. We mixed HeLa-CD81-Snorkel-tag EVs enriched by ultrafiltration with human platelet EV concentrates from three different donors pre-cleaned by differential centrifugation to remove platelets and large EVs. StEVAC purified EVs from mixed populations show similar concentrations compared to HeLa-Snorkel-tag EV elutes (Fig 5A) and the elutes showed HeLa EV specific surface proteins CD44, CD146 and MCSP1 and depletion of platelet EV markers CD31, CD41b, CD42a, CD62P, CD40 and CD49e (Fig 5B; Fig S5A). Next, we mixed human plasma EVs isolated by differential ultracentrifugation with three individual HeLa-Snorkel-tag EV concentrates enriched by ultrafiltration. Indeed, NTA results showed no significant differences in particle concentration in elutes from control and plasma mixed EVs (Fig 5C). MBFCM showed enrichment for HeLa EV specific surface proteins and depletion of plasma EV markers CD31, CD41b, CD42a, CD62P, CD69 and HLA-DRDPDQ^22,23^ (Fig 5D; Fig S5B). In addition, StEVAC purified EVs are actively taken up into Huh-7 cells at similar efficiency as freshly enriched EVs (Fig 5E; Fig S6). This suggests that Snorkel-tag EVs can be isolated from complex sample matrices, paving the ground for better understanding EV cargo under various conditions and - if inserted into conditional, transgenic mice - even in a tissue or cell dependent manner ex vivo.

**Figure 5:**
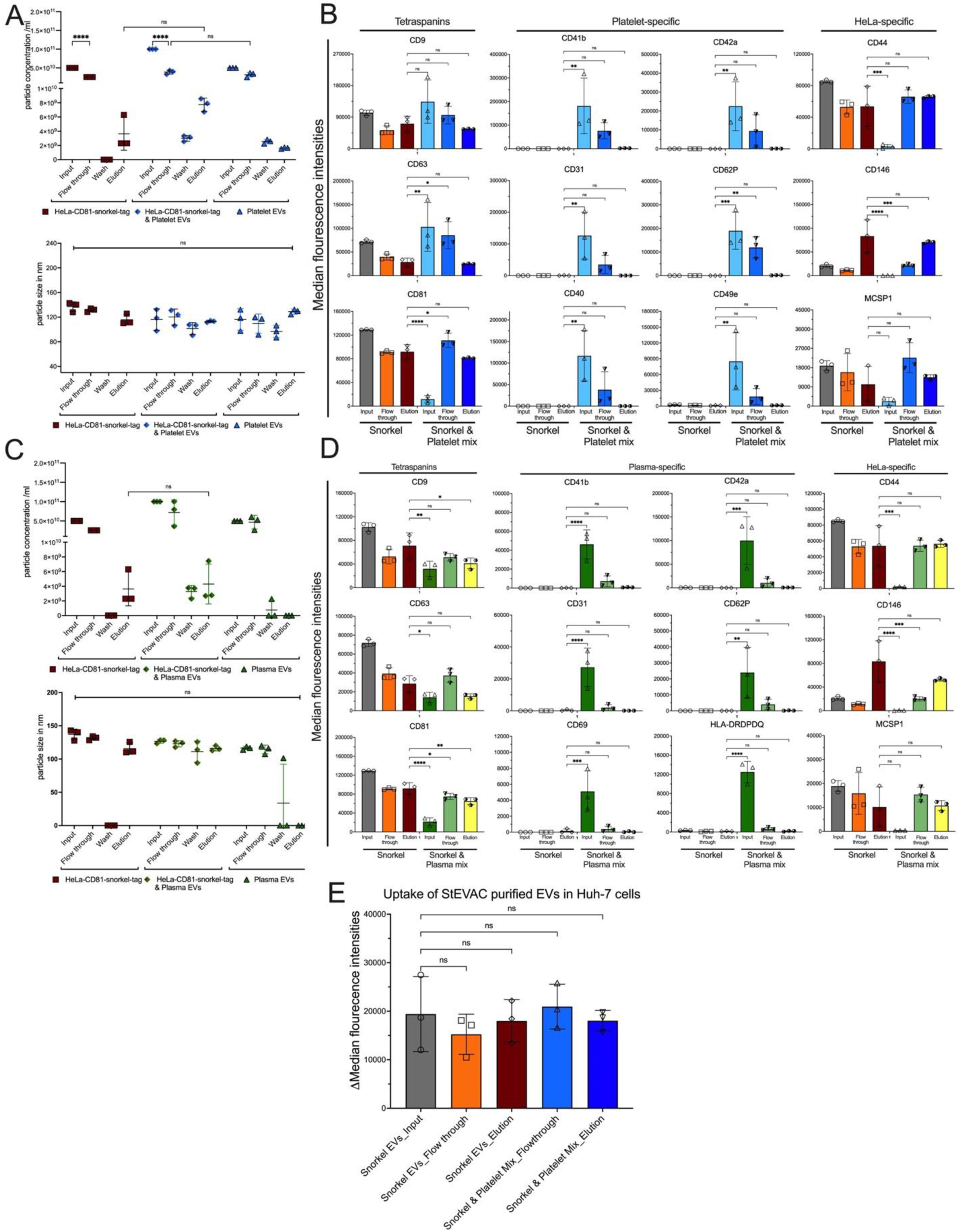
Confirming StEVAC for purification of EVs from complex matrices. StEVAC purification and characterization of HeLa-CD81 Snorkel-tag EVs from human platelet EV enriched mix. **A,** Nanoparticle tracking analysis (NTA) counts of purified EVs from HeLa-CD81-Snorkel-tag and mix of human platelet EVs (n= 3 individual donors), eluted after incubation with PreScission protease treatment overnight and a following wash step. **B,** Multiplex bead-based flow cytometry (MBFCM) assay results for input, flowthrough and elution of EVs from HeLa-CD81-Snorkel-tag and HeLa-CD81-Snorkel-tag mixed with human-platelet EVs. CD44, CD146 and MCSP1 are enriched only in the elutes for HeLa-CD81-Snorkel-tag EVs (right column), platelet EV specific CD31, CD41b, CD42a, CD62P, CD40 and CD49e are depleted (center 2 columns). StEVAC purification and characterization of HeLa-CD81 Snorkel-tag EVs from human plasma mix. **C,** Nanoparticle tracking analysis (NTA) counts of purified EVs from HeLa-CD81-Snorkel-tag and mix of human platelet EVs, eluted after incubation with PreScission protease treatment overnight and a following wash step (n=3). **D,** Multiplex bead-based flow cytometry (MBFCM) assay results for input, flowthrough and elution of EVs from HeLa-CD81-Snorkel-tag and HeLa-CD81-Snorkel-tag mixed with human-plasma EVs. CD44, CD146 and MCSP1 are enriched only in the elutions for HeLa-CD81-Snorkel-tag EVs (right column), plasma EV specific CD31, CD41b, CD42a, CD62P, CD69 and HLA-DRDPDQ are de-riched (center 2 columns). Tetraspanins CD9, CD63 & CD81 are enriched equally in both the elutions. **E,** Flowcytometric analysis of uptake StEVAC purified EVs from mixture of platelet concentrates labelled with CLIP-505 substrate into Huh-7 cells. Unpurified EVs (or) input EVs and flowthrough EVs were used as control (n=3). Same number of particles were used for controls and elutes in uptake. One-way ANOVA was applied on values; nsP>0,05, *P < 0.05, **P < 0.01, ***P < 0.001. 2-way ANOVA multiple comparison test was used for analysis of (A) and (C). 1-way ANOVA multiple comparison test was used for analysis of (B) and (D). Data are shown as mean+_SD, *P < 0.05, **P < 0.01, ***P < 0.001.

### Snorkel-tag EV miRNA cargo is highly similar to wild type EVs

In addition to HeLa cells, as an example of a therapeutically relevant EV origin, CD81 Snorkel-tag construct was introduced into telomerized MSCs, WJ-MSC/TERT273 cells, by lentiviral transduction and selected for CD81 Snorkel-tag expression. Stably expressing WJ-MSC/TERT273 cells showed typical specific markers CD73, CD90 and CD105 (data not shown). Conditioned media from WJ-MSC/TERT273 WT and CD81 Snorkel-tag cells were concentrated by 100 MWCO Amicon filters and StEVAC was performed. NTA results demonstrate ∼30 % of Snorkel-tag EVs in the elute (Fig 6A). However, we observed a slight background signal from the WT elutes with no significant differences in diameter of the particles (Fig 6B). To evaluate the purity of elutes, we performed bead-based flow cytometry by staining CD9, CD63, CD81 and FLAG on flow compatible CD81 magnetic beads. In contrast to NTA results, we observed strong signals for tetraspanins and FLAG in Snorkel-tag elutes and no signals in the elutes from WT EVs (Fig 6C). This results strongly suggests the specificity of StEVAC also in the context of MSC-EVs.

**Figure 6.**
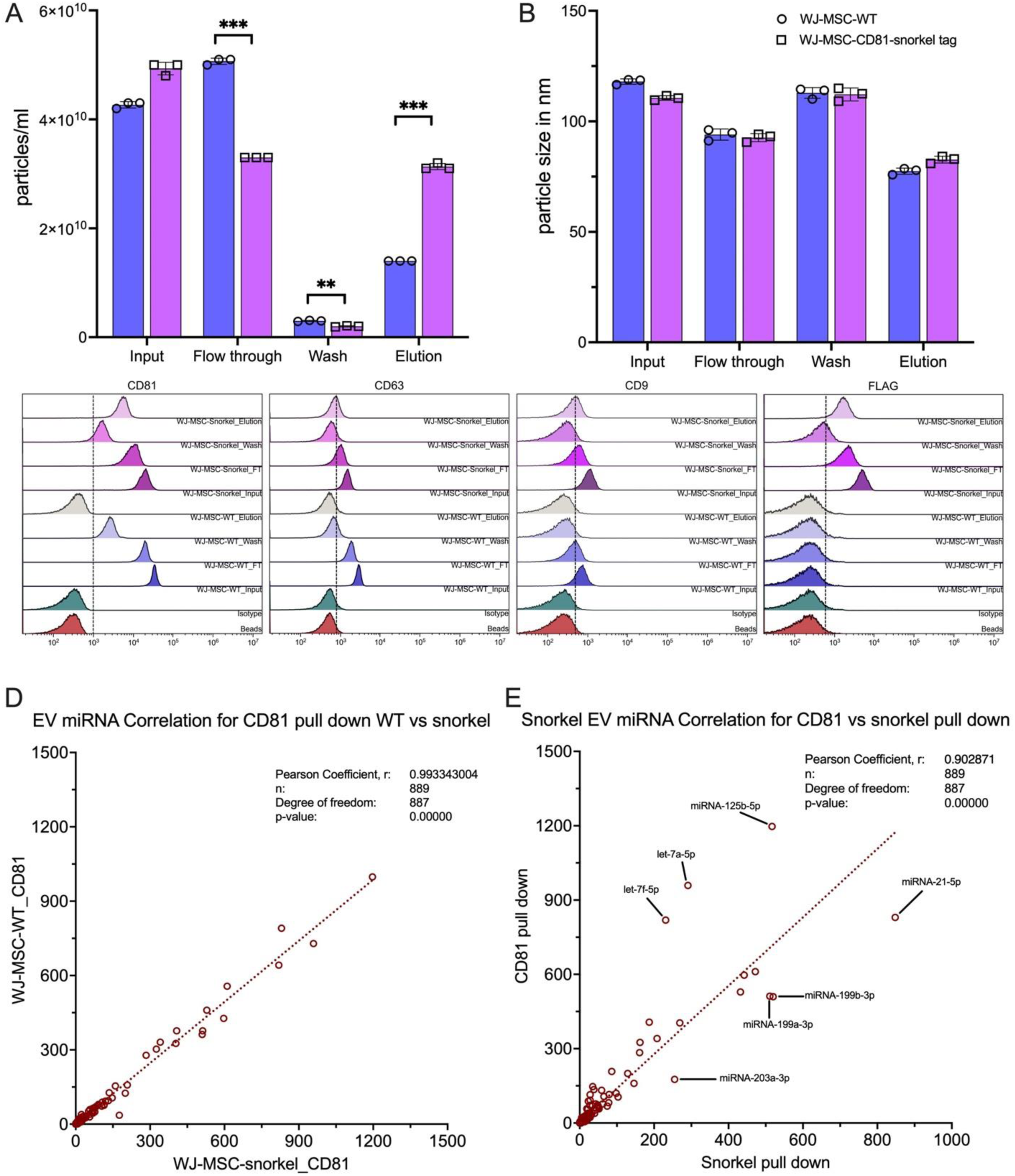
Influence of Snorkel-tag on EV cargo loading. StEVAC purification of EVs from WT and CD81-Snorkel-tag WJ-MSCs. **A,** Nanoparticle analysis (NTA) shows significant differences in elutes of WT versus CD81-Snorkel-tag EVs. **B,** no significant differences were observed in size distribution between flowthrough, washes and elution compared to input. **C,** Bead-based flow cytometry evaluation of flow throughs, washes and elutes from WJ-MSC-WT and WJ-MSC-CD81-Snorkel-tag EVs for CD63, FLAG-tag and CD9. **D,** Pearson correlation scatter plots of miRNAs expression levels between WJ-MSC derived EVs isolated by anti-CD81 pull-down from CD81-Snorkel-tag enriched EVs and WT EVs (n=3). **E,** Pearson correlation scatter plots of miRNAs expression levels between WJ-MSC derived CD81-Snorkel-tag enriched EVs purified by either CD81 specific pull-down or by Snorkel-tag pull down shows only minor differences in cargo loading (n=3).

To elucidate if overexpression of CD81-Snorkel-tag would alter the miRNA pattern associated with MSC-EVs, small RNA sequencing of EVs immunoprecipitated by anti-CD81 antibodies from WJ-MSC-CD81 Snorkel-tag versus wild type cells was performed using 3 independent purifications each per group. A total of 889 miRNAs were identified in all samples with an almost perfect correlation (Pearson r^2^=0.99) and no miRNA differences reaching statistical significance (Fig 6D, Fig S7C). Similarly, we compared the miRNA cargo of Snorkel-tagged EVs immunoprecipitated by anti-HA versus anti-CD81 antibodies resulting in a very high correlation (Pearson r^2^=0.90), with only 7 out of 889 (0.78 %) miRNAs being significantly differentially present (Fig 5E, Fig S7D), an overview of total reads from nine libraries is shown in Fig S7A. Out of approximately 20 million reads in any sample analyzed, miRNAs showed less than 10% enrichment in EVs (Fig S7B). All other, more abundant RNA species, including mRNA, rRNA, tRNA, snoRNA, and snRNA were filtered from the total raw reads. These results suggest only minimal differences induced by stable Snorkel-tag overexpression in EVs.

## Discussion

Previously reported EV subtypes and recent advances in nanoparticle isolation and analytical methods have led to identification of ever-increasing EV subtypes and non-vesicular extracellular nanoparticles (NVEPs) such as lipoproteins, ribonucleoproteins and recently discovered exomeres^24,25^ and supermeres^26–28^ increasing the complexity of their already heterogenous nature. Recently, MISEV 2023, an expert consensus position paper has taken a pragmatic approach for EV nomenclature by classifying EVs <200 nm as small EVs (sEVs) or >200 nm as large EVs (lEVs) even irrespective of their biogenesis^29^.

Considering the EV complexity and heterogeneity involved, purification strategies capable to isolate single EV species will play a pivotal role in defining their final functionality. Over the last decades, various EV isolation methods have been developed, each with its advantages and limitations. Differential ultracentrifugation (dUC) remain as the most commonly used EV enrichment method which provides relatively high purity but comes with major pitfall due to limitation in scalability and co-precipitation of protein aggregates^30,31^. Ultrafiltration and TFF are alternatives offering less harsh conditions as compared to dUC and are better scalable^32^. Size exclusion chromatography has emerged as a leading method to achieve EVs without co-contaminants from biological fluids or when combined with dUC or TFF^33^, however it is unclear at this point, if this also removes bioactive EV molecules^34,35^. Immunoprecipitation is another alternative method for EV isolation from biological fluids. However, it can be costly and the choices of antibodies may bias the isolation to certain EV subtypes^36^, while elution after capture is difficult without destroying the integrity of EVs.Taken together, EV heterogeneity, co-isolation of contaminants, EV integrity and cargo preservation, functional assay compatibility, standardization and reproducibility are major bottlenecks in understanding of EV-based therapeutics^37^. Similarly, standards of specific Evs are necessary to define their intrinsic heterogeneity, source and recovery efficiency in EV separation methods^38,39^.

Recent reports on heparin-^40,41^, tim4-^42^, aptamer-^43^ based affinity purifications showed increase in purity of EVs, however, detailed investigations on EV functionality and co-isolation of glycoproteins are needed^40,41,44^. Additionally, use of high salts or chelating agents for elution can influence on EV membrane integrity and functionality.

In this study, we developed a Snorkel-tag based extracellular vesicle affinity chromatography (StEVAC) which allows on-column isolation of EVs in non-destructive way. CD81, tetraspanin family member, is the most highly enriched EV membrane protein in almost all subtypes of EVs^45^. On cell surfaces, CD81 plays an important role in adaptive immunity by its interaction with CD19 and CD21 in B cell receptor signaling^46^. It has been shown that mutations in CD81 resulted in disruption of CD19 complex formations on B cell leading to antibody deficiency syndrome in humans^41^. Additionally, transmembrane domain (TM1) and large extracellular loop (LEL) of CD81 play an important role in trafficking, localization of co-receptors and signaling^46–51^. Considering the wide spread presence of CD81 on various EVs, we genetically fused Snorkel-tag, specially designed for transmembrane proteins such as GPCRs, ion-channels where both N- and C-termini of protein are inside the cell. This also enables that the Snorkel-tag is separated from CD81 by linker sequences and does not interfere with CD81 functionality. However, we point out, that we did not investigate if signaling functions due to the protein engineering would have been disturbed, as we focused on using the genetically encoded CD81-Snorkel-tag fusion protein that displays the Snorkel-tag components on the surface of EV membranes to allow for easy isolation and covalent labeling of EVs in vitro.

We thus systematically evaluated StEVAC for purifying EVs harboring the Snorkel-tag from cell culture supernatants of by spiking them into blood derived EVs followed by rigorous characterizations. Therefore, we mixed Snorkel-tag harboring EVs with complex matrices such as plasma EVs and platelet concentrates, and still purified them without non-Snorkel-tagged EV contaminants. This will pave the way to generate transgenic animals, from which EVs can be isolated in a cell or tissue and time resolved manner for assessing not only cargo but also functionality due to the mild elution conditions.

To harness the therapeutic potential of EVs it is a pre-requisite to reliably track the biodistribution of EVs in vivo in a quantifiable way for clinical applications. Several pre-clinical studies have addressed the pharmacokinetics and biodistribution of EVs by directly labelling EVs with fluorescent lipophilic dyes such as DiR, DiD and PKH26/67^52–54^. However, fluorescent lipid dyes have longer half-life and are reported to form nano-sized micelles resulting in inaccurate spaciotemporal detection of EVs^55,56^. Fluorescent and bioluminescent reporters such as PalmtdTomato^55^, nanoluciferase^57^, mCherry^58^, palmGRET^59^ provide an alternative when genetically fused to EV enriched proteins or lipids. However, the use of fluorescent reporters is limited due to high background and low tissue penetration of signals^60,61^. On the other hand, radioactive tracers such as technetium hexamethylpropylene amine oxime (^99m^Tc-HMPAO)^62^, iodine-131^63^, indium-111-oxine (^111^In)^64^ and ^89^Zirconium deferoxamine ([^89^Zr]Zr-DFO)^65^ have been applied to label EVs for pharmacokinetics and biodistribution analysis using single-photon emission computed tomography (SPECT), positron emission computed tomography (PET). However, use of radioisotopes comes with limitations as it requires certified facilities and well-trained operators, making these techniques costly. To overcome these limitations, we introduced the CLIP-tag as a component of the Snorkel-tag, which is a covalent self-labeling protein-tag that can be specifically and irreversibly labeled with O^2^-benzyl-cytosine (BC-) derivatives with the advantage that the majority of model organisms does not react with benzyl-cytosine derivatives, avoiding any endogenous activity and background^66^. Furthermore, a number of commercially available cell-permeable and impermeable BC fluorophores can be used not only in vitro or after EV purification, but also for labeling of EVs in tissue samples or histology sections after delivering the EVs to avoid the dyes to alter the biodistribution. Still, even if Snorkel-tagged EVs behave similarly in vitro-uptake experiments, it cannot be excluded, that the Snorkel might impact their in vivo distribution.

Our results in any case demonstrate that labeling of Snorkel-tag in cells or on EVs after StEVAC method is fast, and allow tracking of EVs upon uptake. It has been shown that similar self-labeling tag, SNAP-tag, Snap^CaaX^ reporter mice provide a great tool for in vivo live labeling and imaging of tissues^67^.

While tetraspanin-based reporters such as exomap1^68^ and CD63-EGFP^69^ generated transgenic animal models have been used previously, these reporter constructs contain the transgene inside of the EVs, thus lack the possibilities of using them as affinity isolation means, but most probably do not alter the surface characteristics of these EVs. On the other hand, CD63^FLAG^-EGFP^70^ and truncated CD9-EGFP^71^ transgenic mouse models enable tracking and affinity purification of EVs, but the extracellular domains of the tetraspanins are altered, which can produce unintended biochemical changes in tetraspanins altering their functionality, post-translational modifications, subcellular localizations. Since the Snorkel-tag is placed after a 5th transmembrane domain distant from unaltered CD81 domains, it might be less likely to interfere with natural CD81 functionality but still is displayed to the outside, allowing not only tracking but also isolation to advance our knowledge on EV biogenesis, cell-type specific, physiologic and pathologic function and cargo in or ex vivo. Still, it is necessary to carefully characterize the recombinant EVs for potential changes, several of which were covered here showing no major influence of our genetically engineered CD81. Taken together, this technology platform might contribute to a detailed understanding of EV biology and advance our development of EV based therapeutics and diagnostics.

## Materials and Methods

### Cell culture

Cell culture experiments were performed under sterile and antibiotic free conditions. Human dermal fibroblasts (HDFs) from adult skin three healthy donor (HDF76), and WJ-MSC/TERT273 were provided by Evercyte GmbH. Cells were grown in DMEM/Ham’s F-12 (1:1 mixture) (BIOCHROME, Germany) supplemented with 10 % fetal calf serum (FCS) and 4 mM L-Glutamine (Sigma Aldrich GmbH St Louis, MO, USA) at 7% CO2 and 37°C. WJ-MSCs were cultivated in Mesencult media with supplements (Stem cell) at 7% CO2 and 37°C. HEK293 cells were cultivated in DMEM with Na-pyruvate (BIOCHROME, Germany) supplemented with 10 % fetal calf serum (FCS) and 4 mM L-Glutamine (Sigma Aldrich GmbH St Louis, MO, USA) at 7% CO2 and 37°C. Huh-7 cells were provided by Samir EL Andaloussi Department of laboratory medicine, Karolinska Institutet, Stockholm, Sweden. Huh-7 cells were cultivated in DMEM (ThermoFisher Scientific) supplemented with 10% fetal calf serum (FCS) (Sigma Aldrich GmbH St Louis, MO, USA) and 1 X GlutaMAX (ThermoFisher Scientific) at 5% CO2 and 37°C. HeLa cells were grown in RPMI 1640 (ThermoFisher Scientific) supplemented with 10% fetal calf serum (FCS) (Sigma Aldrich GmbH St Louis, MO, USA) and 1 x GlutaMAX (ThermoFisher Scientific) at 5% CO2 and 37°C.

### Generation of stable cell lines

For CD81-Snorkel-tag, insert was cloned into pLVX-IRES-Hygro vector. Lenti-X^TM^ 293T cells (TaKaRa; Cat.no 632180) were transfected with pLVX-IRES-Hygro-CD81-Snorkel-tag plasmid using Lenti-X packaging single shots according to manufacturer instructions. In brief, Approximately, 24 hr before transfection, 4 x 10^6^ Lenti-X 293 cells were seeded in T75 flask in 8 ml growth media and incubated at 37° C, 5% CO2 overnight. 7 μg of pLVX-IRES-Hygro-CD81 snorkel-tag vector plasmid DNA was diluted to 600 μl with sterile nuclease free water. Diluted plasmid vector DNA was added to Lenti-X packaging single shots (TaKaRa; Cat.no: 631276) and vortexed for 20 sec at high speed. Transfection mixture was incubated at room temperature for 10 min to allow nanoparticle complexes to form. After 10 min incubation, 600 μl of nanoparticle complex was added dropwise to 8 ml of cell culture prepared. Incubate the cells at 37° C, 5% CO2 overnight, add additional 6 ml of fresh growth medium and incubate at 37°C, %CO2 for additional 48 hr. After 48 hr incubation, harvest the lentiviral supernatants and filter through 0.45 μm filter. Transfer clarified supernatant to fresh container and combine 1 volume of Lenti-X concentrator (TaKaRa; Cat.no: 631232) to 3 volumes of clarified supernatant. Mix by gentle inversion and incubate overnight ar 4°C. Post incubation, centrifuge sample at 1500 x g for 45 min at 4° and pellet was suspended gently in 1/10^th^ of original volume of growth media. 100 μl aliquots of virus stocks were prepared and stored in −80°C for future use. For generating stable cells, target cells (HeLa and/or WJ-MSC-TERT 293 cells) were seeded in T75 flasks 18 hr before transduction. Virus stock thawed and mixed with growth medium containing polybrene (4 μg/ml), viral supernatant with polybrene was added to cells and transduced for 16 hrs. After transduction, virus containing media was replaced with normal growth media and culture for another 24 hr. After 24 hr of recovery, medium was replaced with 20 μg/ml hygromycin containing fresh growth media and selected for live cells after 4-7 days. Once stable cells were generated, they were characterized and prepared master cell bank for future use.

### Human platelet concentrates & plasma

Single donor platelet concentrates were provided by the Red Cross Blood Transfusion Service (Linz, Upper Austria). All samples were collected during routine thrombocyte apheresis in accordance with the policies of the Red Cross Transfusion Service, Linz. All blood donors signed their informed consent that residual blood material can be used for research and development purposes. All experimental protocols were approved by and carried out in collaboration with the Red Cross Blood Transfusion Service, Linz.

Platelet concentrates were generated by single platelet apheresis using an automated cell separator (Trima Accel Automated Blood Collection System, TerumoBCT) at the Red Cross Blood Transfusion Service (Linz, Upper Austria). All blood donors signed an informed consent that blood material can be used for research and the study was conducted in accordance with the policies of the Red Cross Transfusion Service. Platelets were finally stored in SSP+ (Macopharma) and ACD-A (acid citrate dextrose + adenosine, Haemonetics® anticoagulant citrate dextrose solution, Haemonetics®, Braintree) was used as an anticoagulant. 2 mL of the platelet concentrate (containing ∼1 × 10^6^ platelets/µL) were aseptically transferred into a separate storage bag and experiments were carried out within 24 h after donation.

Plasma was provided by the Red Cross Blood Transfusion Service (Linz, Upper Austria). All samples were collected collected in accordance with the policies of the Red Cross Transfusion Service, Linz. All blood donors signed their informed consent that residual blood material can be used for research and development purposes. All experimental protocols were approved by and carried out in collaboration with the Red Cross Blood Transfusion Service, Linz.

### EV isolation procedures

#### Tangential flow filtration

Conditioned media from HeLa-WT and HeLa-CD81-Snorkel-tag overexpressing cells were collected and subjected to a low speed spin at 700 × g for 5 minutes at 4° C to remove cellular debris, followed by 2000 × g spin for 10 minutes at 4°C to remove larger particles and cell debris. The supernatant was then sterile filtered with a 0.22 μm filter cups. Conditioned media was diafiltrated using 2 volumes of initial volume to ∼ 35 ml using KR2i TFF system (SpectrumLabs) with 300 kDa cut-off hollow fibre filters (MidiKros, 370 cm2 surface area, SpectrumLabs) at a flow rate of 100 ml/min (transmembrane pressure at 3.0 psi and shear rate at 3700 sec−1. Diafiltrate was further concentrated to ∼1 ml using Amicon ultra-15 centrifugal filter unit (Catalog # UFC910024) at 4°C with 3500 x g. Concentrated EV solution was quantified for size and concentration and 100 µl aliquotes were stored in PBS-HAT buffer^10^ at −80°C for subsequent characterization studies.

#### Ultrafiltration

Pre-cleaned conditioned media (700 x g for 5 min and 2000 x g for 10 min) from HeLa-WT and HeLa-CD81-Snorkel-tag overexpressing cells was sterile filtered using syringe (VWR) with cellulose acetate membrane filters (0.22 µm pore size) to remove any larger particles. The filtered conditioned media was ultrafiltrated using 100 kDa MWCO Amicon ultra-15 cenrifugal filter unit (Catalog # UFC910024) at 4°C with 3500 x g (Balaj et al. 2015). The concentrate was diafiltrated with 2 volumn of 1 x PBS and concentrated to final volume of ∼1 ml. Final volume was quantified by NTA for size and concentration of EVs. After quantification EV sample were freshly used for further purification by StEVAC method.

#### Ultracentrifugation

Human plasma EVs were isolated by ultracentrifugation as follow: 250 ml of fresh citrate plasma frozen at −25°C was provided by the Red Cross Blood Transfusion Service (Linz, Upper Austria, Austria). Plasma was thawed on ice, centrifuged twice at 2500 g for 10 min and 5 ml of plasma was diluted 1:5 with PBS and rest of the plasma were aliquoted with 10 ml volumes and stored at −80° C for future experiments. 1:5 diluted plasma was centrifuged at 14,000 g for 30 min at 4°C. After centrifugation, supernatants were centrifuged at 100,000 g at 4°C for 90 min (Eppendorf). Supernatants were separated and pellets were suspended in 1 ml of centrifuged supernatants to enrich for plasma EVs in plasma matrices^16,17^.

For human platelet EV enrichment, fresh platelets from all the donors were incubated for 2 hr on a shaker at room temperature^15^. After incubation, samples were diluted 1:1. Platelets and cell debris were removed with a centrifugation for 5 min at 5,000 g, 14,000 g for 30 min followed by 0.2 μm filtration. Filtered samples were immediately used for NTA, StEVAC and MBFCM.

### Nanoparticle Tracking Analysis (NTA)

Nanoparticle tracking analysis was applied to determine particle size and concentration of all samples. All samples concentrated by TFF and UF were characterized by NTA with a NanoSight NS500 instrument equipped with NTA 2.3 analytical software and an additional 488 nm laser. Samples were diluted to 1:1000 in sterile filtered PBS (0.22 µm filter). Diluted samples were loaded in the sample chamber with camera level 13. Four to five 30 sec videos were recorded per sample in light scatter mode with 5 sec delays between each recording. Screen gain 10, detection threshold 7 were kept constant for all the recordings. Using batch process facility all the measurements were analyzed automatically.

### Snorkel-tag based Extracellular Vesicle Affinity Chromatography (StEVAC)

Conditioned media from HeLa-WT and HeLa-CD81-Snorkel-tag overexpressing cells were processed using ultrafiltration-based EV isolation. Isolated EVs were quantified using NTA and ∼2.5 x 10^10^/ml were incubated with 250 ml of anti-HA magnetic beads (Catalog # 88836, ThermoFisher Scientific; bead concentration 10 mg/ml) overnight at 4° C on a rotospin test tube rotator. Post incubation, beads were separated on magnetic rack and unbound EV solution was collected and beads were washed with 0.22 mm filtered PBS. After washing step, beads were suspended in 0.22 mm filtered PBS with 5 ml (10 units) of PreScission protease (catalog # 27084301; GE healthcare Life Sciences) and incubated overnight at 4°C on a rotospin test tube rotator for on-column PreScission protease cleavage. After overnight incubation, tubes were placed on magnetic rack for separating beads and elutes were collected. Collected elutes along with flow through and wash samples were quantified for size and concentration using NTA. For characterization studies samples were freshly used. For pre-mixed samples with HDF derived EVs, ∼2.5 x 10^10^/ml particles were with equal number of HeLa-CD81 Snorkel-tag enriched particles as inputs. For human platelet and plasma ∼5 x101^0^/ml particles from human-platelet or from plasma were mixed with HeLa-CD81 Snorkel-tag EVs as inputs.

### Immunoblotting

HeLa-WT cells and HeLa-CD81-Snorkel-tag expressing cells were collected and the cell pellet was lysed with 100 µL of RIPA buffer, kept on ice, and vortexed five times every 5 min. The cell lysate was then spun at 12,000 × g for 10 min at 4°C and the supernatant was transferred to a new tube and kept on ice. Protein concentrations for the supernatants were quantified by BCA assay (ThermoFisher Scientific) according to manufacturer’s instructions. 50 µg of cell lysates and 1 x 10^9^ to 5 x 10^9^ particles were mixed with buffer containing 0.5 M dithiothreitol, 0.4 M sodium carbonate (Na2CO3), 8% SDS, and 10% glycerol, and heated at 95°C for 10 min. The samples were loaded onto a NuPAGE Novex 4–12% Bis-Tris Protein Gel (Invitrogen, Thermo Fisher Scientific) and run at 120 V in NuPAGE MES SDS running buffer (Invitrogen, Thermo Fisher Scientific) for 2 h. The proteins on the gel were transferred to an iBlot nitrocellulose membrane (Invitrogen, Thermo Fisher Scientific) for 7 min using the iBlot system. The membrane was blocked with Odyssey blocking buffer (LI-COR) for 1 hour at room temperature with gentle shaking. After blocking, the membrane was incubated overnight at 4°C or 1 hour at room temperature with primary antibody solution (CD81, Santa Cruz Biotechnology, sc-166029, 1:200; TSG101, abcam, ab125011, 1:1,000; Alix, abcam, ab117600, 1:2,000; Syntenin-1, Origene, TA504796, 1:1,000; Calnexin, abcam, ab22595; HA-tag, Cell Signaling, 3724, 1:1,000; SNAP/CLIP-tag, NEB, P9310S, 1:1,000; FLAG-tag, Sigma, F3165, 1:5,000). The membrane was washed with PBS supplemented with 0.1% Tween-20 (PBS-T, Sigma) three times for every 5 min and incubated with the corresponding secondary antibody (LI-COR; anti-mouse IgG, 926-68072, 1:10,000; anti-rabbit IgG, 925-32213, 1:10,000) for 1 hour at room temperature. Finally, the membrane was washed with PBS-T for three times with 5 min interval, twice with PBS and visualized on the Odyssey infrared imaging system (LI-COR) at 700 and 800 nm.

### CLIP-tag labeling quantification by Flow cytometry

HeLa-WT cells and HeLa-CD81-Snorkel-tag overexpressing cells were collected and suspended in 1 ml of growth media (RPMI 1640 + 10% FCS + 1 X GlutaMAX). To this, non-cell-permeable CLIP-substrate (CLIP-SurfaceTM 647; Catalog #S9234, NEB) with final dilution of 1:100,000 was added and incubated at for 1 hour at 95% humidity, 5% CO2 and 37°C. Post incubation, cells were spun at 300 x g for 5 min to remove the unlabeled dye and washed twice with PBS and pelleted at 300 x g for 5 min and resuspended in 100 ml of PBS. Dead cells were excluded by 4’,6-diamidino-2-phenylindole (DAPI) staining and doublets were excluded by forward/side scatter area versus height gating. Samples were kept on ice and measured with MACSQuant Analyzer 10 flow cytometer (Miltenyi Biotec). GraphPadPrism 8.2.1 (GraphPadPrism Software, La Jolla, CA, USA) was used to analyze data and assemble figures.

### Multiplex bead-based flow cytometry assay for EV surface protein profiling

Different sample types were subjected to bead-based multiplex EV analysis by flow cytometry (MACSPlex EV IO Kit, human, Miltenyi Biotec). Unless indicated otherwise, EV-containing samples were processed as follows: Samples were diluted with MACSPlex buffer (MPB) to, or used undiluted at, a final volume of 60 µL and loaded onto wells of a pre-wet and drained MACSPlex 96-well 0.22 µm filter plate before 3 µl of MACSPlex Exosome Capture Beads (containing 39 different antibody-coated bead subsets) were added to each well. Generally, particle counts quantified by NTA, and not protein amount, were used to estimate input EV amounts. We used 1 x 10^9^ particles as input EV amounts. Filter plates were then incubated on an orbital shaker overnight (14–16 hours) at 450 rpm at room temperature protected from light. To wash the beads, 200 µl of MPB was added to each well and the filter plate was put on a vacuum manifold with vacuum applied (Sigma-Aldrich, Supelco PlatePrep; −100 mBar) until all wells were drained. For counterstaining of EVs bound by capture beads with detection antibodies, 135 µl of MPB and 5 µl of each APC-conjugated detection antibody cocktail (anti-CD9, anti-CD63, and anti-CD81) were added to each wells and plates were incubated on an orbital shaker at 450 rpm protected from light for 1 h at room temperature. Next, plates were washed by adding 200 µL MPB to each well followed by draining on a vacuum manifold. This was followed by another washing step with 200 µl of MPB, incubation on an orbital shaker at 450 rpm protected from light for 15 min at room temperature and draining all wells again on a vacuum manifold. Subsequently, 150 µl of MPB was added to each well, beads were resuspended by pipetting and transferred to V-bottom 96-well microtiter plate (Thermo Scientific). Flow cytometric analysis was performed using MACSQuant Analyzer 10 flow cytometer (Miltenyi Biotec). All samples were automatically mixed immediately before 70–100 µl were loaded to and acquired by the instrument, resulting in approximately 3,000–5,000 single bead events being recorded per well. FlowJo software (v10, FlowJo LLC) was used to analyze flow cytometric data. Median fluorescence intensity (MFI) for all 39 capture bead subsets were background corrected by subtracting respective MFI values from matched non-EV buffer (PBS) that were treated exactly like EV-containing samples (buffer/medium + capture beads + antibodies)^21^. GraphPadPrism 8.2.1 (GraphPadPrism Software, La Jolla, CA, USA) was used to analyze data and assemble figures.

For indirect labeling of Snorkel-tag, samples were incubated with capture beads overnight (14-16 hours) at 450 rpm at room temperature protected from light. The beads were washed with 200 µl MPB and 135 µl MPB added to each well and 15 µl of anti-HA-tag antibody (1:1000 final dilution) was added and incubated for 1 hour at room temperature on an orbital shaker at 450 rpm protected from light. After incubation, the MPB in the wells was drained and wash steps were repeated. For counterstaining of anti-HA-tag antibody labeled on EVs bound capture beads, 135 µl of MPB and 15 µl of Dylight-649 conjugated to anti-rabbit detection antibody (1:1000 final dilution) was added and plates were incubated on an orbital shaker at 450 rpm protected from light for 1 hour at room temperature. After incubation, MPB was drained, washed and flow cytometry analysis was performed using MACSQuant Analyzer 10 flow cytometer (Miltenyi Biotec) as mentioned above. MFI for all 39 capture bead subsets were background corrected by subtracting respective MFI values from matched non-EV buffer (PBS) that were treated exactly like EV-containing samples (buffer + capture beads + Dylight 649 detection antibody).

MBFCM EV quantification for flow throughs and elutes in pre-mixed human platelet EV and plasma EVs, 5 x 10^9^ EVs/ ml for inputs, 2.5 x10^9^ EVs/ml for flow throughs and 2.5 x 10^8^ EVs/ ml for elutes were subjected to bead-based multiplex EV analysis by flow cytometry. Rest of protocol was performed according to manufacturer instructions in tube format. Flow-cytometric analysis was performed using CytoFLEX S (Beckman Coulter). Data was analysed with CytExpert.

### Flow cytometry of bead-bound extracellular vesicles

For staining of surface EV tetraspanins and Snorkel-tag, flow throughs and washes from StEVAC method were suspended in 20 µl of Exosome-human CD81 flow detection beads (Thermo Scientific, 10622D) and incubated at 4°C overnight while rotating. The next day, bead bound EVs were separated by placing on a magnetic rack. Bead-bound EVs were suspended in 100 µl of PBS with antibody cocktail (CD63 Antibody, anti-human, FITC, REAfinity™, 130-118-076; CD9 Antibody, anti-human, APC-vio770, REAfinity™, 130-118-813; anti-FLAG, APC, REAfinity™, 130-119-584) incubated at 4°C for 1 hr while rotating. After staining, bead-bound EVs were washed twice with PBS and suspended in 300 µl of PBS. Samples were kept on ice and data was acquired at a CytoFLEX S (Beckman Coulter). Data was analysed with CytExpert. Median fluorescence intensity was normalized over the isotype controls (ΔMFI).

### Fluorescence microscopy for cells

HeLa-WT and HeLa-CD81-Snorkel-tag overexpressing cells were seeded onto coverslips or μ-slides (ibidi GmbH, Martinsried, Germany) and incubated over night at 37°C. Following day, cells were fixed with 4% paraformaldehyde for 15 min, washed two times with PBS, and permeabilized for 10 min in 0.3% Triton X-100 followed by two PBS-washes. Cells were blocked with 2% BSA for 30-60 min. After blocking, slides were incubated in primary (HA-tag, Cell Signaling, 3724, 1:800; CD81, Thermo Scientific, 11525542, 1:500) and secondary antibody (AF488, AF647, Jackson immunoResearch, 1:1,000) solutions prepared in 2% BSA solution for 60 and 30 min respectively in a humidified chamber at room temperature, each followed by 3 washes in PBS. Hoechst 33342 was included for counterstaining of DNA right before the last wash step.

For staining non-permeabilized cells, no fixation and permeabilization steps were involved. Cells were blocked in 2% BSA for 60 min and were directly incubated with primary and secondary antibodies as mentioned above.

### Cellular uptake of StEVAC purified EVs

For quantification of cellular uptake of StEVAC purified EVs, equal number of eluted EVs from CD81-Snorkel-tag, WT and CD81-Snorkel-tag input EVs were labelled with CLIP-substrate (CLIP-Surface^TM^ 647; Catalog #S9234, NEB and/or CLIP-Surface^TM^ 488; Catalog #S9232, NEB) with final dilution of 1:100,000 was added and incubated at for 1 hour at 95% humidity, 5% CO2 and 37°C. Post-incubation, unlabeled dye was removed by desalting columns. Labelled EVs were added to human hepatocellular carcinoma cells (Huh-7) seeded a day before at density of 3 x 10^4^ cells per well in a 96-well plate. Cells were incubated for 2 hrs at 95% humidity, 5% CO2 and 37°C.

#### For flow cytometry

After incubation, cells were washed twice with ice cold PBS, typsinzed, spun down at 900 x g for 5 min and resuspended in 100 μL of PBS. Dead cells were excluded from analysis via 4′-,6-diamidino-2-phenylindole (DAPI) staining and doublets were excluded by forward/side scatter area versus height gating. Samples were kept on ice and data was acquired at a CytoFLEX S (Beckman Coulter). Data was analysed with CytExpert. Mean fluorescence intensity was normalized over the control/untreated cell sample (ΔMFI)

#### For fluorescence microscopy

After incubation, cells were washed twice with ice cold PBS. Hoechst33342 staining for nuclear counter staining. Next, cells were stained with lysotacker^®^ Green DND-26 at 50 nM concentrations and analyzed immediately with Leica TCS SP8.

### Transmission Electron Microscopy (TEM)

For TEM analysis we used 4 different protocols based on the sample and application.

All the solutions used for the staining procedure were pre-filtered using 0.22 μm filter units (Milipore/VWR).

Freshly prepared EVs were adhered on Athene Old 300 mesh copper grids (Agar Scientific, Stansted, Essex, UK) and fixed with 1% glutaraldehyde. Grids were washed three times with nuclease free water (NFW) and stained for 5 min with 2% phosphotungstic acid hydrate (Carl Roth, Karlsruhe, Germany). The grids were left to dry and the specimens were visualized using TEM (FEI Tecnai T20, FEI Eindhoven, Netherlands) operated at 160 kV.

For all other TEM images, 5 µl of sample were added onto glow-discharged formvar-carbon type B coated electron microscopy grids for 3 min. Samples were removed by using wet whatmann filter paper. Grids were either prepared for immunogold labeling (see below) or carefully washed twice with filtered PBS. After washes, 5 µL of filtered 2% uranyl acetate were added for 10-30 sec, uranyl acetate was removed using wet whatman filter paper, grids were air dried and using visualized using a transmission electron microscope (Tencai 10).

For immunogold labeling, grids were blocked after the initial binding step of the sample using filtered 2% BSA (in PBS) for 10 min. Primary and secondary antibodies were diluted in 0.2% BSA solution. After blocking, grids were placed on 15 µL primary antibody solution (anti-CD81, 1:50; anti-HA-tag, 1:50) for 60 min. Post incubation, grids were washed with 0.2% BSA 6 times and incubated with secondary antibody (goat anti-mouse secondary antibody conjugated with 10 nm gold particles & goat anti-rabbit secondary antibody conjugated with 4 nm gold particle) with dilution of 1:50 for 60 min. After incubation, grids were washed 6 times with PBS followed by 6 washing steps with ddH2O. Finally, grids were stained with 0.2% uranyl acetate for 10 to 30 sec. Excess uranyl acetate was removed using a wet whatman filter paper, grids air dried and visualized using a transmission electron microscope (Tencai 10).

### EV RNA isolation

Freshly enriched EVs from WJ-MSC WT and CD81-Snorkel-tag cell supernatants were immunoprecipitated either with hCD81 exosome isolation kit (Miltenyi biotec; 130-110-914) or µMACS™ HA Isolation Kit (Miltenyi biotec; 130-091-122). Trizol LS (Thermo Scientific) was added to EV precipitates. RNA was extracted by miRNeasy Mini Kit (Qiagen; 217004) and enriched for small RNA by RNeasy MinElute Cleanup Kit (Qiagen; 74204) according to manufacturer instructions. The RNA integrity was tested on the Agilent 2100 Bioanalyzer using smallRNA kits (Agilent Technologies). RNA concentrations were calculated by using the Bioanalyzer software. For small RNAs this was calculated for all lengths spaning the chip (5-200 bp).

### Library preparation and RNAseq

EV RNA samples were prepared for small RNA sequencing using QIAseq small RNA Library Prep kit (Qiagen), which includes the adapter ligation at the two ends of small RNAs, reverse transcription and amplification. During the reverse transcription, the unique molecular identifier (UMI) was added on each molecule. The quality of finished libraries were checked on Agilent 2100 Bioanalyzer using High sensitivity DNA chip and the quantity of libraries were measured by qPCR using KAPA Library Quantification kit (Roche). Libraries were pooled with equal amount and sequenced on an Illumina Novaseq sequencer by single-end 100 bp sequencing.

### Bioinformatic analysis

The raw data was quality filtered and trimmed by fastx_toolkit, and adaptor sequences were removed using Cutadapt. The reads were collapsed to remove identical UMI-containing reads. FastQC was used to ensure high-quality sequencing data. The tRNA reads were filtered and the rest reads were mapped to miRNAs from miRBase v22 using Bowtie allowing zero mismatches but allowing for non-templated 3′ A and T bases. The small RNA expression profiles generated were used for differential expression analysis in R using the DESeq2 package, and volcano plots were generated using R.

### Statistical analysis

Statistics were either calculated with Excel or Graph Pad Prism, and respective tests are indicated below figures in result sections. ± Standard deviations were derived from at least 3 independent experiments. Two tailed tests were performed using an error probability of 0.05. If not indicated, the experiments were performed less than three times.

## Data Resources

The accession number for the RNA-seq reported in this paper is GEO: GSE251842.

## Acknowledgements

This project has been supported by international PhD programme “BioToP-Biomolecular Technology of Proteins”, funded by the Austrian Science Fund (FWF). In addition, this project was supported by the BOKU Core Facilities for Biomolecular and Cellular Analysis and Multimodal Imaging. Authors also thank the Eppendorf for their generous support with instrumentation.

## Disclosure statement

Johannes Grillari and Regina Grillari are co-founder and shareholder of Evercyte GmbH. Matthias Hackl serves as a CEO and co-founder, Johannes Grillari and Regina Grillari are co-founders and serves on the scientific advisory board of TAmiRNA GmbH. André Görgens, Dhanu Gupta and

Samir EL Andaloussi are consultants for and have equity interests in Evox Therapeutics Ltd., Oxford, United Kingdom. All other authors declare no potential conflict of interests.

## Authors contribution

Conceptualization, M.R.B, A.G, J.K, S.E.A, J.G; Methodology, M.R.B, A.G, Y.Y, M.H, J.J, M.S, R.G; Experiments performed by M.R.B, A.G, Y.Y, S.V, D.G, G.C, S.B, C.P, S.W, M.P, D.G, G.C, S.B, S.V; M.R.B wrote the manuscript; J.K, S.E.A and J.G supervised the project.

## Notes

https://www.ncbi.nlm.nih.gov/geo/query/acc.cgi?acc=GSE251842

