## Supplementary data for "Snorkel-tag Based Affinity Chromatography for Recombinant Extracellular Vesicle Purification"

[12]Austrian Cluster for Tissue Regeneration

Corresponding author at: Ludwig Boltzmann Institute for Traumatology. The Research Center in Cooperation with AUVA, Vienna Austria

### Supplementary notes

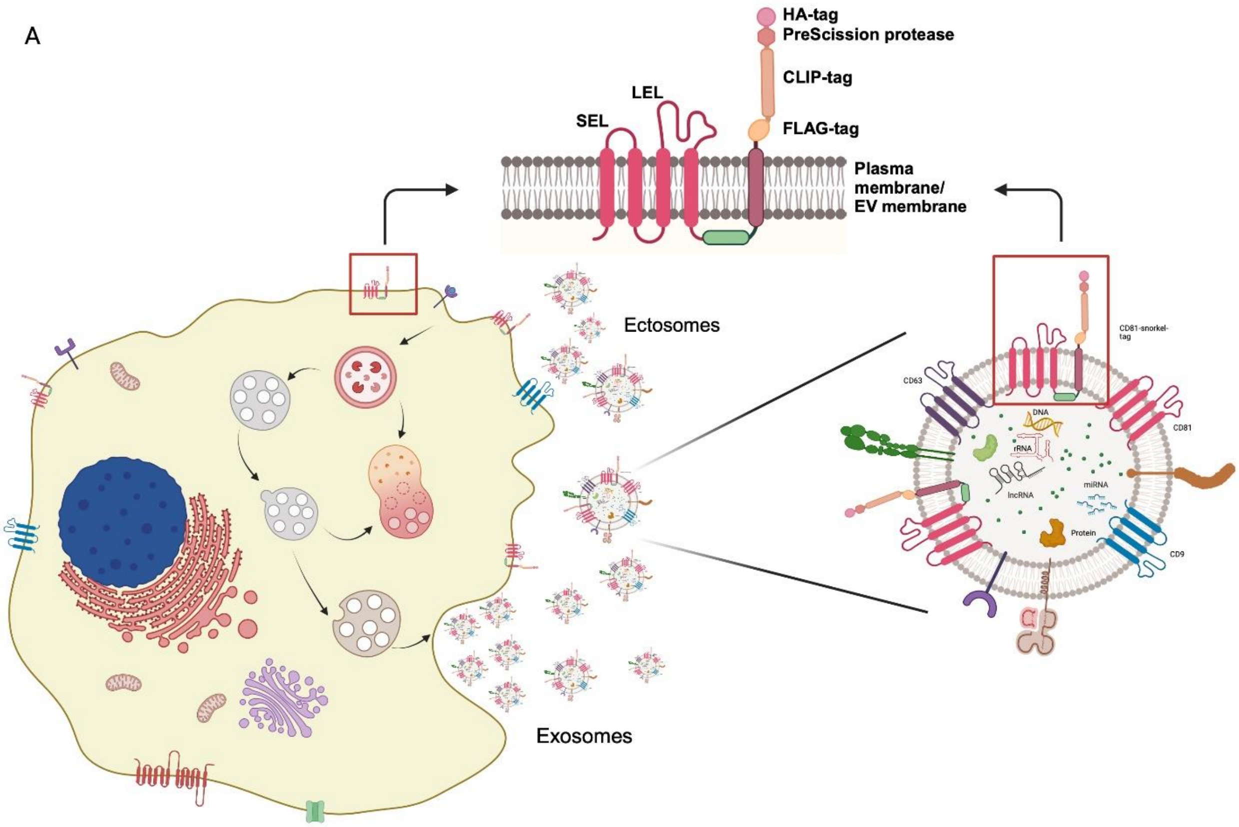

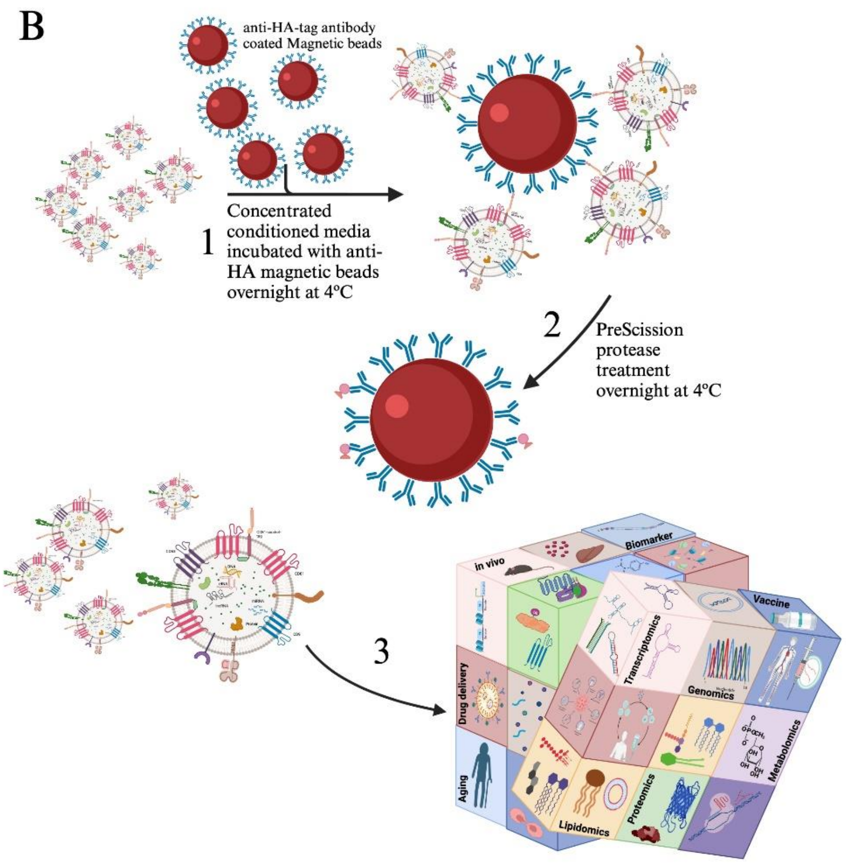


(A) Schematic representation of the Snorkel-tag genetically fused to the C terminus of CD81on cell and EV surface membranes. (B) Overview on Snorkel-tag based EV affinity chromatography (StEVAC): (1) Concentrated conditioned media from CD81-snorkel-tag stably expressing cells, incubated overnight with anti-HA matrix at 4ºC to capture snorkel-tag enriched EVs. (2) Mild treatment of captured EVs with PreScission protease at 4ºC releases EVs without changing their biophysical properties. (3) StEVAC method enables to understand the cargo of EVs under normal and disease physiological states, when expressed under tissue-specific promoter in transgeneic mouse models

Supplementary figure 1

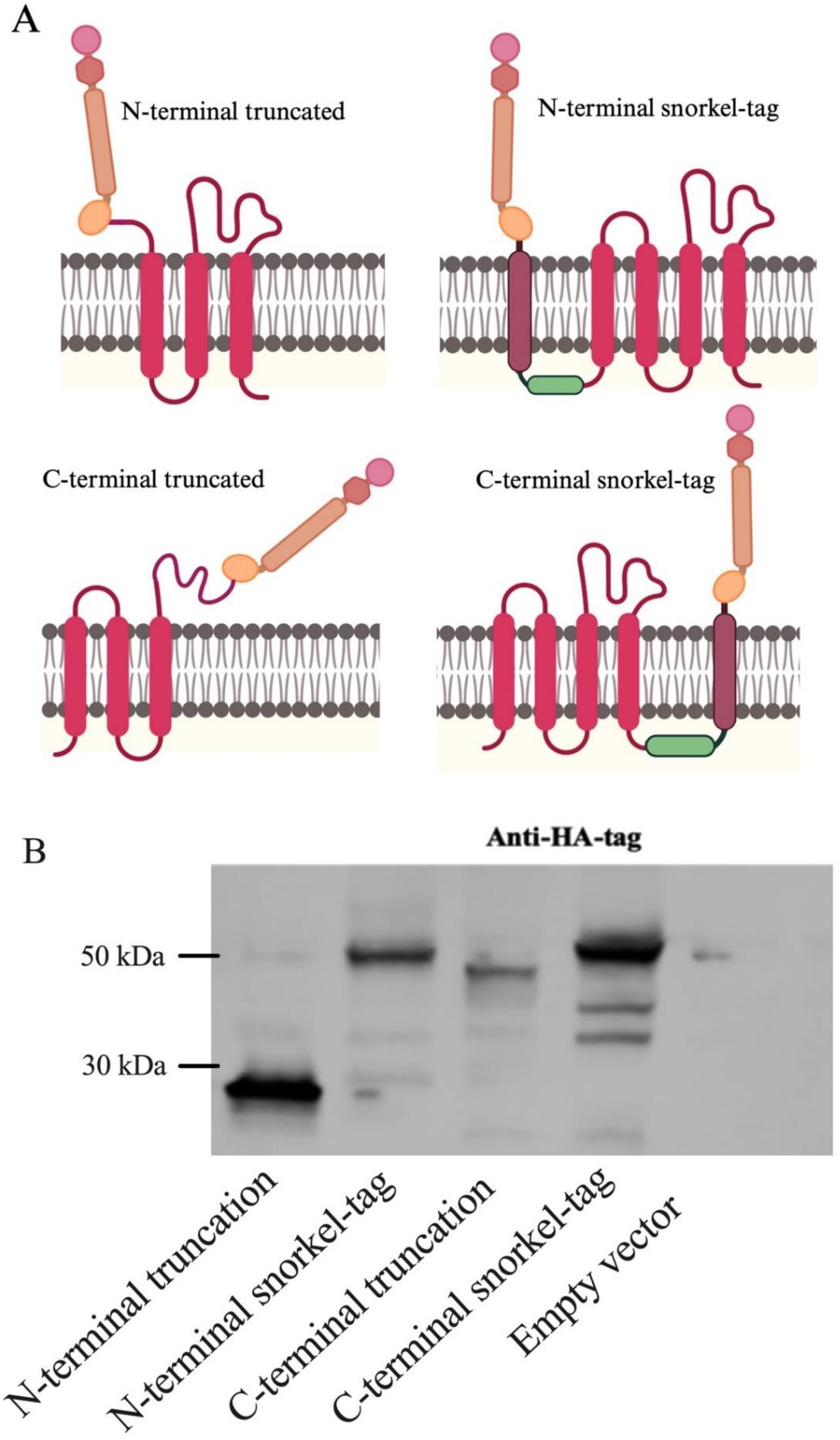

**Figure S1** (A) Schematic representation of full length CD81 genetically fused with snorkel-tag at C-termini and CD81 truncated versions devoid of either transmembrane domain 1 or 4 with snorkel-tag fused to SEL or LEL respectively. (B) Western blot of anti-HA tag for all four fusion proteins transiently expressed in HeLa cells. Western blot results reveal N-terminal truncated version of CD81 snorkel-tag did not express full length protein.  Created with BioRender.com.

#### Supplementary figure 2

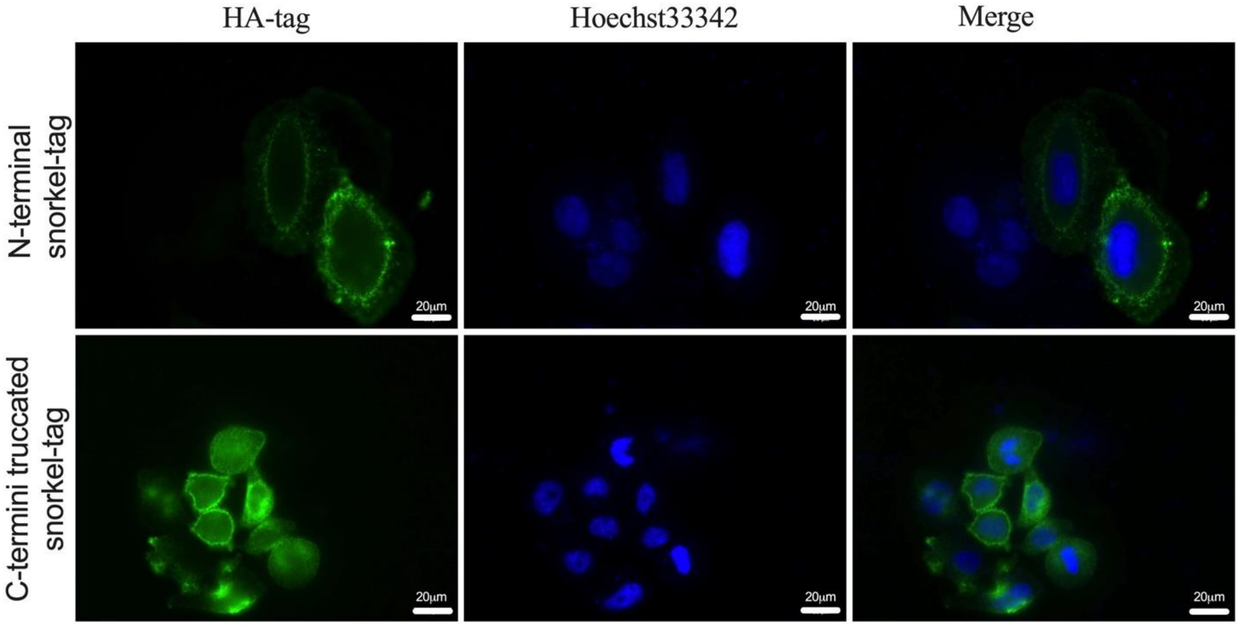


**Figure S2** Fluorescent images of fixed HeLa cells expressing CD81 with N-terminal snorkel-tag and C-terminal truncated snorkel-tag stained with anti-HA tag antibody and Alexafluor-488 anti-rabbit secondary antibody

### Supplementary figure 3

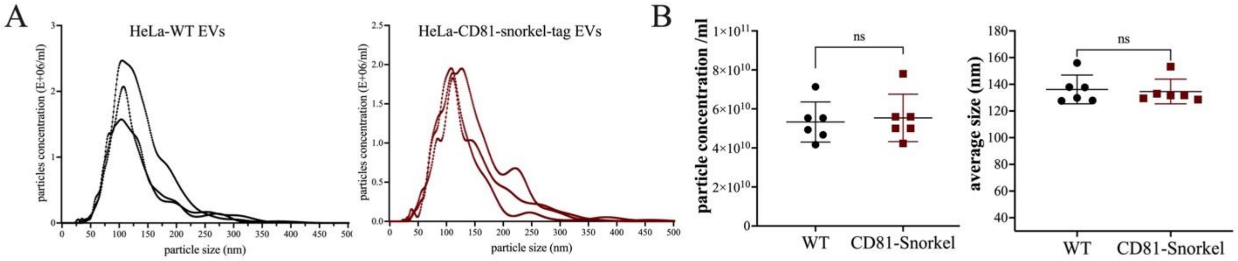


**Figure S3 A,** Representative particle size and concentration for EVs derived from HeLa-WT and HeLa-CD81-snorkel-tag cell lines (n=3). **B,** Particle concentrations and size of ultrafiltrated particles from 75 ml conditioned media from 6 individual experiments. Unpaired t-test was applied on raw values; nsP > 0.05.

### Supplementary figure 4

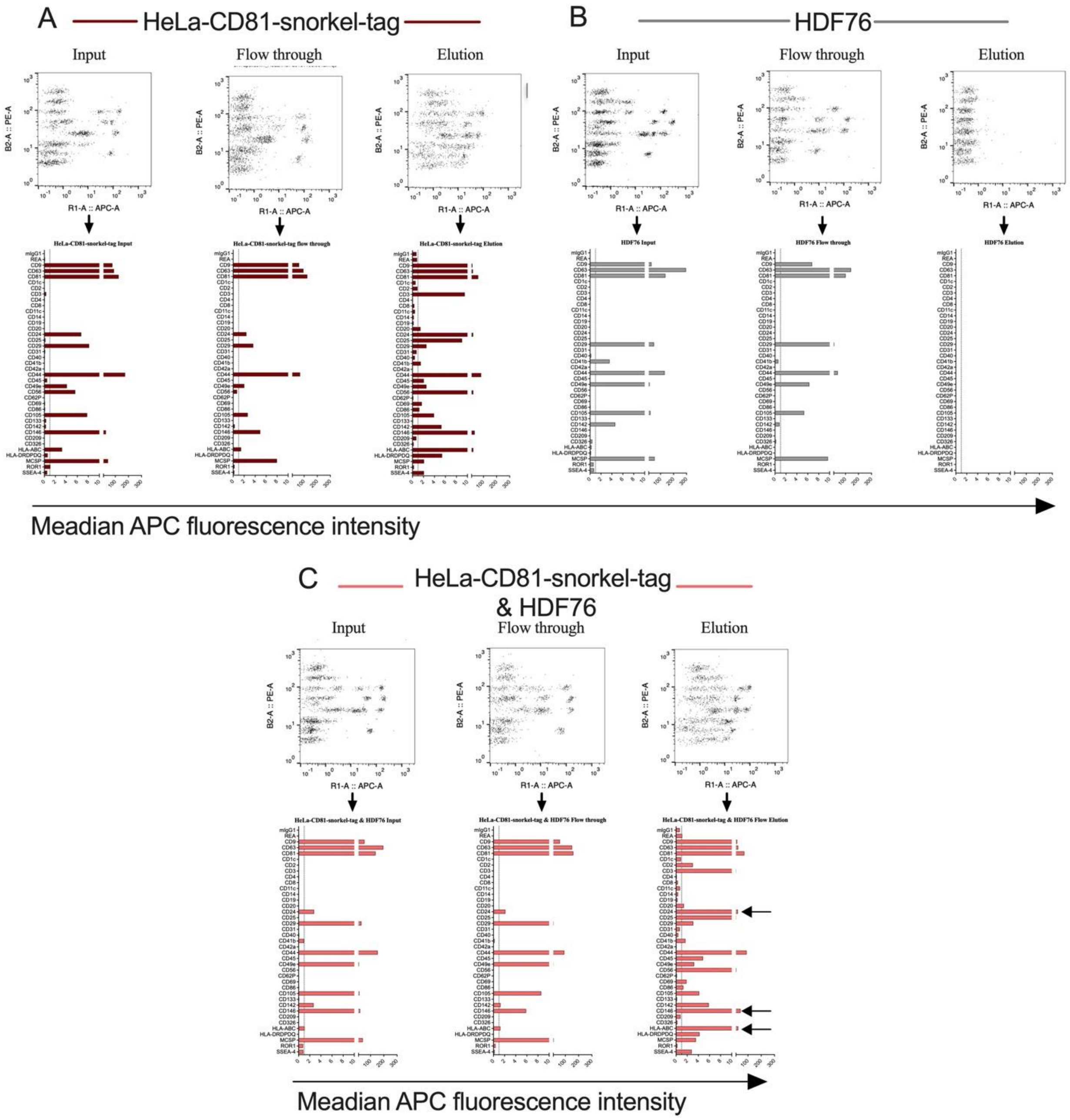


**Figure S4** Confirming StEVAC to purify EVs carrying snorkel-tag from mixed population of EVs. Multiplex bead-based assay results for input, flowthrough and elution of EVs from HeLa-CD81-snorkel-tag **A**, HDF76 **B** and HeLa- CD81-snorkel-tag mixed with HDF76 **C**.

Supplementary figure 5

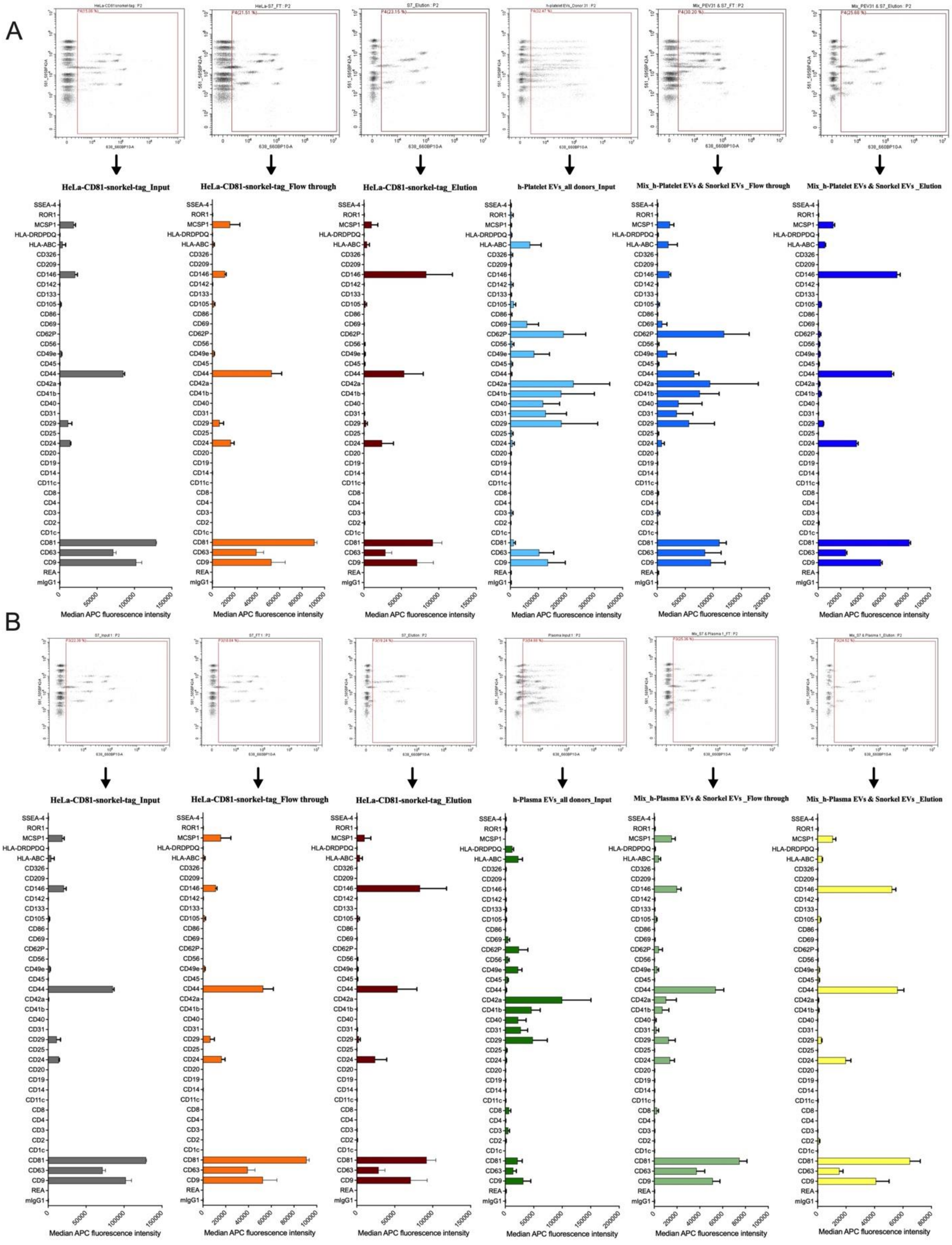

**Figure S6** MACSPlex 37 EV surface protein panel for HeLa-CD81-Snorkel tag and in mixtures with human platelet (A) and with human plasma (B); inputs, flowthroughs and elutes probed by anti-pan tetraspanin APC antibodies (n=3).

#### Supplementary figure 6

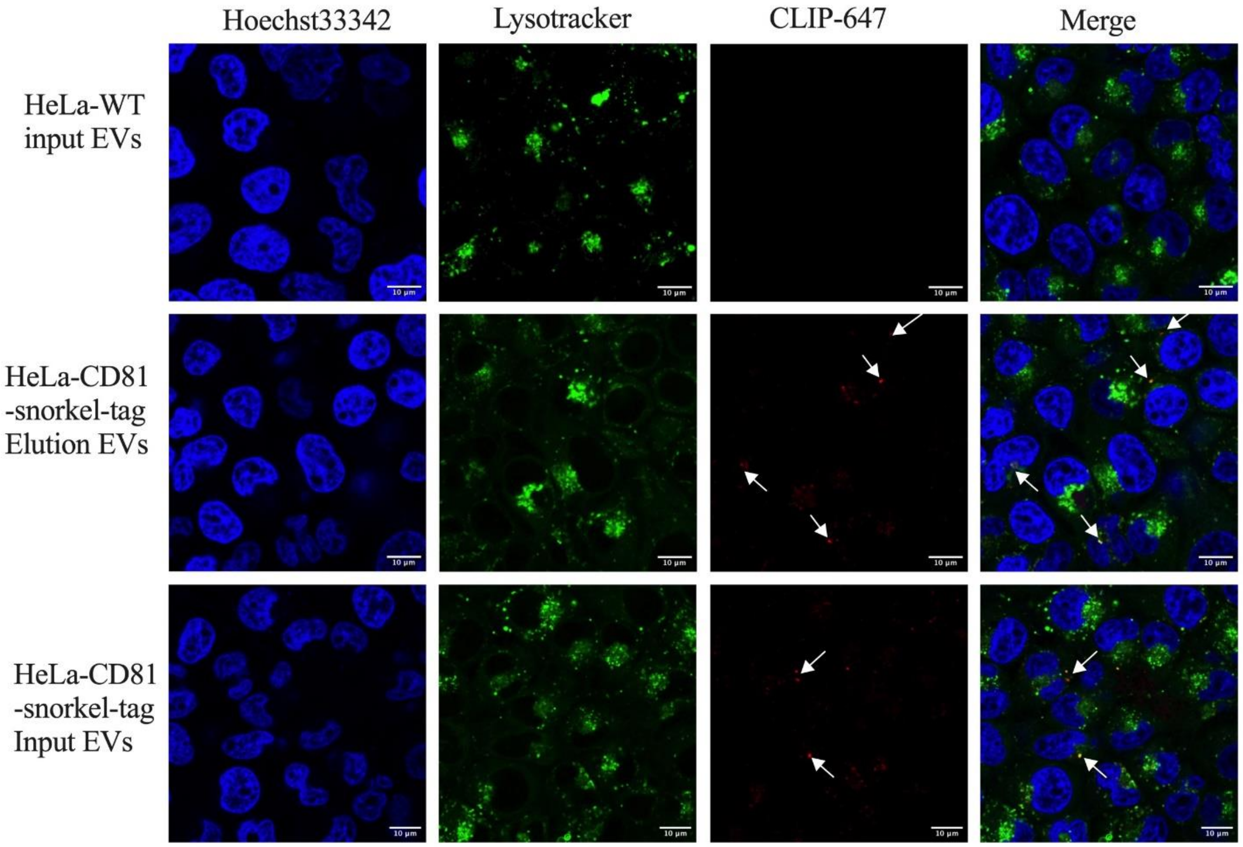


**Figure S6** Uptake of StEVAC purified EVs in Huh-7 recipient cells. Representative Confocal images of StEVAC purified EVs labelled with CLIP-647 uptake in Huh-7 cells from HeLa-WT, HeLa-CD81-snorkel-tag, as a positive control HeLa-CD81-snorkel-tag unpurified EVs in red. Counter staining with LysoTracker in green.

Supplementary figure 7

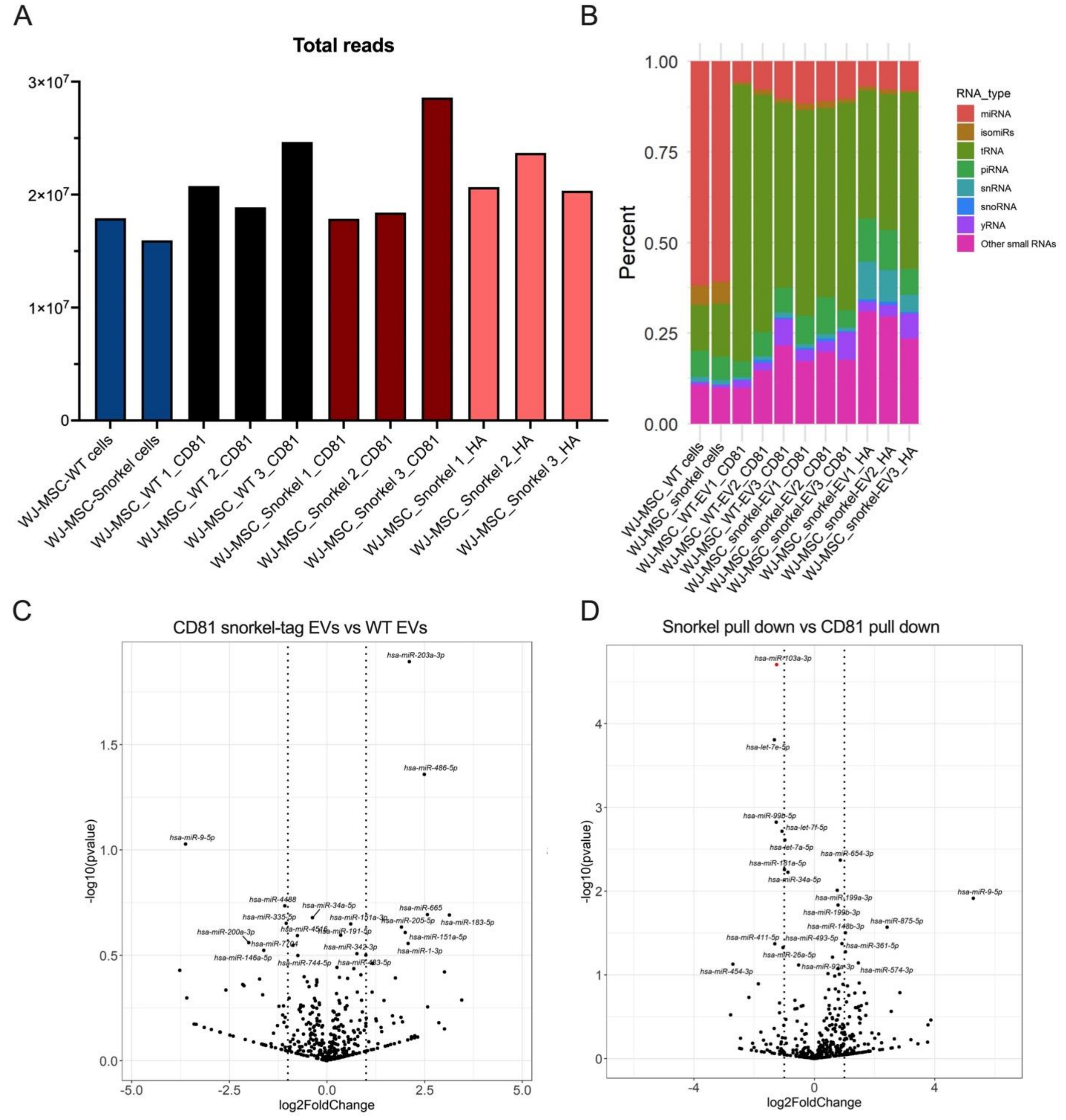


**Figure S7** EV cargo characterization **A,** total reads of small RNA sequencing from WJ-MSC WT and snorkel-tag EVs pull down from snorkel-tag and CD81. **B,** percentage of small RNA species enriched in EVs. **C,** Volcano plot shows no significant differences in miRNAs between EVs immunoprecipitated by anti-CD81 antibodies from WJ-MSC-CD81 snorkel tag and wildtype. **D,** Volcano plot identifying single miRNA differentially expressed in snorkel tag enriched EVs immunoprecipitated by anti-HA and anti-CD81 antibodies.
